# Language impairment with a microduplication in 1q42.3q43

**DOI:** 10.1101/2020.05.26.117903

**Authors:** Antonio Benítez-Burraco, Maite Fernández-Urquiza, Mª Salud Jiménez-Romero

**Author notes:** Corresponding author: Antonio Benítez-Burraco, Área de Lingüística General. Departamento de Lengua Española, Lingüística y Teoría de la Literatura. Facultad de Filología. Universidad de Sevilla. C/ Palos de la Frontera s/n. 41007-Sevilla (España).

## Abstract

Deletions and duplications of the distal region of the long arm of chromosome 1 are associated with brain abnormalities and developmental delay. Because duplications are less frequent than deletions, no detailed account of the cognitive profile of the affected people is available, particularly, regarding their language (dis)abilities. In this paper we report on the cognitive and language features of a girl with one of the smallest interstitial duplications ever described in this region, affecting to 1q42.3q43 (arr[hg19] 1q42.3q43(235,963,632-236,972,276)x3). Standardized tests as well as the analysis of her language use in natural settings suggest that the proband’s speech is severely impaired, exhibiting dysarthric-like features, with speech problems also resulting from a phonological deficit boiling down to a verbal auditory memory deficit. Lexical and grammatical knowledge are also impaired, impacting negatively on both expressive and receptive abilities, seemingly as a consequence of the phonological deficit. Still, her pragmatic abilities seem to be significantly spared, granting her a good command on the principles governing conversational exchanges. In silico analyses (literature mining, network analysis) and in vitro analyses (microarray) point to several genes as potential candidates for the observed deficits in the language domain. These include one gene within the duplicated region (*LYST*), one predicted functional partner (*CMIP*), and three genes outside the 1q42.3q43 region, which are all highly expressed in the cerebellum: *DDIT4* and *SLC29A1*, found strongly downregulated in the proband compared to their healthy parents, and *CNTNAP3*, found strongly upregulated.

## INTRODUCTION

Deletions and duplications of the distal region of the long arm of chromosome 1 result in developmental anomalies, including intellectual disability and language problems. Deletions usually entail microcephaly, seizures, cognitive and psychomotor delay, growth delay, facial dysmorphisms, and cardiac defects (Juberg et al., 1981; Roberts et al., 2004; van Bon et al., 2008). That said, the exact symptoms exhibited by patients depend on the extension of the deleted region and consequently, on the number and functions of the affected genes. Duplications of this region are less frequent and patients exhibit a more variable clinical presentation, although common symptoms include intellectual disability and delayed psychomotor development, facial anomalies, and macrocephaly (Hemming et al., 2016; Morris et al. 2016).

In this paper, we report on a girl with an interstitial duplication in the 1q42.3q43 region. Rearrangements of this region are rare. Silipigni and colleagues (2017) reviewed 7 cases found in the literature, affecting to a slightly broader region (1q42.13q43), and described 2 new cases: a deletion of 1q42.13q43 and the reciprocal duplication, found in two consanguineous probands. Subjects with duplications suffer from language deficits. Specifically, the child examined by Silipigni et al. (2017) showed cognitive and social delay, and said his first words at the age of 2 only. Nonetheless, a detailed characterization of the language problems exhibited by patients with duplications of the distal region of the long arm of chromosome 1 is still pending. Not surprisingly, no conclusive genotype-phenotype link has been found for language deficits. One reason is that the role of the duplicated genes in neurodevelopment, and particularly, in language development, is mostly unknown. A second reason is that patients usually bear complex translocations involving other chromosomes. In this paper we provide a comprehensive account of the language deficits and strengths of our proband, including a detailed psycholinguistic profile and an in-depth analysis of her language in use. We further advance a hypothesis about the molecular causes of language dysfunction. For this, we have relied on available evidence of the role played by the duplicated genes in brain development and function, but also, on predicted functional links between some of these genes and selected strong candidates for language development and language evolution in the species. Additionally, we have relied on results of our analysis of whole-genome gene expression pattern of the blood of our proband compared to her healthy parents.

## MATERIAL AND METHODS

### Linguistic, cognitive, and behavioral assessment

The global developmental profile of the proband was evaluated with the revised version of the Batería de Aptitudes Diferenciales y Generales [Battery for Differential and General Abilities] (BADyG) (Yuste and Yuste, 1998) and with the Spanish version of the Inventory for Client and Agency Planning (ICAP) (Montero, 1996). The proband’s linguistic skills were evaluated with the Spanish versions of the Peabody Picture Vocabulary Test (PPVT-3) (Dunn et al., 2006), the Gardner Receptive One Word Picture Vocabulary Test (ROWPVT) (Gardner, 1987), and the Illinois Test of Psycholinguistic Abilities (ITPA) (Kirk et al., 2009). The Registro Fonológico Inducido test [Induced Phonological Register] (Monfort and Juárez, 1988) was used to achieve a detailed knowledge of the proband’s phonological awareness.

#### The Battery for Differential and General Abilities (BADyG)

The BADyG evaluates the child’s abilities in 9 different domains: verbal comprehension (understanding of direct commands involving spatial concepts), verbal reasoning (finding analogical relationships between concepts), numerical comprehension (performing simple arithmetic calculations), numerical reasoning (understanding and solving simple mathematical problems), processing of rotated figures (rotating figures mentally), spatial reasoning (inducing the rules governing the assembly of geometric figures), visuo-auditive memory (recalling meanings from a story which is also displayed graphically), writing abilities (absence of dyslexic-like features), and discrimination (finding subtle differences between geometric figures). Scores are subsequently grouped to obtain 5 composite measures: verbal abilities (verbal comprehension + verbal reasoning), numerical abilities (numerical comprehension + numerical reasoning), visuospatial abilities (processing of rotated figures + spatial reasoning), logical reasoning (verbal reasoning + numerical reasoning + spatial reasoning), and general intelligence (verbal comprehension + verbal reasoning + numerical comprehension + numerical reasoning + processing of rotated figures + spatial reasoning). The version of the test employed in this case was BADyG/E1, aimed for children between 6 and 7 years old.

#### The Inventory for Client and Agency Planning (ICAP)

The ICAP is aimed to evaluate the subject’s functional abilities and maladaptive behaviors in the following general areas: motor skills, social and communication skills, personal living skills, and community living skills. The ICAP measures the frequency and severity of 8 types of behavioral disturbances, which are organized in 3 subscales: asocial maladaptive behavior (uncooperative behavior and socially offensive behavior), internalized maladaptive behavior (withdrawn or inattentive behavior, unusual or repetitive habits, and self-harm), and externalized maladaptive behavior (disruptive behavior, destructive to property, and hurtful to others). Behavior is rated as normal or abnormal, whereas behavioral problems are subsequently rated as marginally serious, moderately serious, serious, or very serious.

#### The Peabody Picture Vocabulary Test (PPVT-3)

This test is aimed to assess the correct acquisition and the level of receptive vocabulary in subject from 2.5 years to 90 years. It consists of 192 sheets with four colored drawings each. After hearing a word, the subject must indicate which illustration best represents its meaning.

#### Gardner Receptive One Word Picture Vocabulary Test (GROWPVT)

This test is aimed to assess receptive vocabulary in children between 2 and 11 years old, as a screening tool for detecting possible speech defects, learning disorders, auditory processing deficits, or problems for auditory-visual association. It consists of 100 picture-naming tasks: after hearing a word, the subject has to choose (either verbally or by pointing) the correct picture out of four different colored pictures.

#### Illinois Test of Psycholinguistic Abilities (ITPA)

This test aims to detect preserved psycholinguistic skills and specific difficulties experienced by children when trying to produce an adequate communicative contribution. In this test, psycholinguistic skills are construed as psychological processes and functions enabling the subject to convey his intentions during the communicative use of language (Kirk and McCarthy, 1961). These psychological skills are ultimately related to the core processes involved in the reception, understanding, and transmission of a linguistic message. The ITPA comprises several subtests that are organized into two levels: the Representative Level and the Automatic Level. The Representative Level assesses the management of symbols via the evaluation of different kinds of processes: i) Receptive processes, including a) the ability to obtain meaning from a spoken word (Listening comprehension), and b) the ability to obtain meaning from a visual symbol, assessed through the capability to choose from among a group of drawings the one that is similar to the stimulus picture (Visual comprehension); ii) Organizational processes, including a) the ability to relate spoken words in a meaningful way, which is tested using a series of verbal analogies of increasing difficulty (Auditory association); and b) the aptitude to relate visual symbols, which is assessed through the matching of the stimulus picture to one drawing chosen among a set of four (Visual association); and iii) Expressive processes, including a) verbal fluency, which is estimated from the number of concepts expressed verbally (Verbal expression), and b) the ability to express concepts through gestures (Motor expression). Regarding the Automatic Level, it assesses how different habits, such as memory or remote learning, are integrated to produce an automatic chain of responses. The assessment is conducted via the evaluation of different kinds of processes: i) Integration or closure tests, including a) the competence to automatically use grammar in a task of sentence completion supported by drawings (Grammar integration or closure), b) the ability to identify a common object from its incomplete representation in a relatively complex context (Visual integration), c) the ability to produce a word from partially pronounced words (Aural integration), and d) the ability to synthesize the separate sounds of a word in order to produce the complete word (Sound gathering); ii) Sequential memory tests, including a) the ability to recognize and produce a word from its partial pronunciation (Auditory sequential memory), and b) the ability to reproduce by heart sequences of non-significant figures after viewing the sequence for a short period of time (Visomotor Sequential Memory).

#### Induced Phonological Register (RFI)

This tests assesses phonological and articulatory abilities of Spanish-speaking children between 3 and 7 years old by means of elicited word production and repetition. Word production is induced by showing the child different pictures that she has to name. If the child is unable to produce the name, the speech-language therapist will provide her with the correct name and will ask the child to repeat it. The test comprises every possible sound of the Spanish language, so that it can be used to detect dyslalia as well as phonological impairment.

#### Language in use

In addition, the proband’s language in use and communication skills were assessed through the analysis of a 12-minutes sample of her talk while chatting with her family. Three different settings were considered: 1) a meal at home with her parents, 2) a family meeting with her parents, her brother and her grandparents; and 3) a conversation with her mother and her brother after one school day. The conversations were video-recorded and then transcribed and coded using the tools provided by the CHILDES Project (MacWhinney, 2000). In brief, speech production phenomena (including relevant information about articulatory processes, prosody, and fluency) were coded in the main lines of the transcript using CHAT (*Codes for the Human Analysis of Transcripts*). Relevant information about gestural communication and the communicative use of gaze was included in the main lines within square brackets. Lastly, phonological, morphological, syntactic, lexical and pragmatic phenomena were coded in dependent lines. Because functional approaches to language impairment demand taking into account the pragmatic consequences of structural errors, we first tagged structural errors and then evaluated their communicative effects using the “Pragmatic Evaluation Protocol of Corpora” (PREP-CORP). This protocol has been satisfactorily used to provide the pragmatic profiles of diverse neurodevelopmental disorders (Fernández-Urquiza et al., 2015, Fernández-Urquiza et al., 2017, Diez-Itza et al., 2018). Finally, a whole pragmatic profile of the proband was built up. For this, we computed the number of occurrences of each label using CLAN (*Computerized Language Analysis*) (Conti-Ramsden, 1996).

#### Assessment of the parents’ cognitive and language abilities

The linguistic profile of the proband’s parents was assessed with the verbal scale of the Spanish version of the WAIS-III (Wechsler, 2002), the Test de Aptitud Verbal “Buenos Aires” [Buenos Aires Verbal Aptitude Test] (BAIRES) (Cortada de Kohan, 2004), the Batería para la Evaluación de los Procesos Lectores en Secundaria y Bachillerato [Battery for the Evaluation of Reading Abilities in High School Students] (PROLEC-SE) (Ramos and Cuetos, 2011), and one specific task aimed to evaluate the comprehension of passive sentences and co-referential expressions. The verbal scale of the WAIS-III comprises 6 tasks aimed to assess the subject’s abilities in two different domains of language processing: verbal comprehension (Similarities (S), Vocabulary (V), Information (I), and Comprehension (CO)) and working memory (Digits (D) and Arithmetic (A)). The BAIRES test aims to evaluate language comprehension and production in subjects above 16 years old. It comprises two tasks which assess, respectively, the subject’s ability to understand language and her ability to provide synonymic words and verbal definitions. Regarding the PROLEC-SE, it is mostly used to assess reading abilities of youngsters between 12 and 18 years old, but because a full reading competence is usually acquired before 18, it can be also used with adults. It comprises six tasks that evaluate three different aspects of reading: lexical processes (word reading, pseudoword reading), semantic aspects (text comprehension, text structure), and syntactic abilities (picture-sentence matching, punctuation marks). To end, the *ad-hoc* task was a sentence-picture matching task aimed to evaluate the comprehension of bound anaphora in complex sentences (that is, the ability to properly bind a noun phrase in the dependent clause to a referential element in the main clause) and of canonical long passive structures (that is, with a *by*-phrase). While the first task is highly-demanding in computational terms, the second task is usually difficult to perform by people with poor reading abilities, because passives are very infrequent in spoken Spanish.

### Cytogenetic and molecular analyses

#### Karyotype analysis

Peripheral venous blood lymphocytes were grown following standard protocols and collected after 72 hours. A moderate resolution G-banding (550 bands) karyotyping by trypsin (Gibco 1x trypsin^®^ and Leishmann stain) was subsequently performed. Microscopic analysis was conducted with a Nikon^®^ eclipse 50i optical microscope and the IKAROS Karyotyping System (MetaSystem^®^ software).

DNA from the patient and her parents was extracted from 100 μl of EDTA-anticoagulated whole blood using MagNA Pure (Roche Diagnostics, West Sussex, UK) and used for subsequent analyses.

#### Fragile X syndrome analysis

CGG expansions affecting the gene *FMR1* (the main determinant factor for X-fragile syndrome) were analyzed in the proband according to standard protocols. Polymerase chain reaction (PCR) of the fragile site was performed with specific primers for the fragile region of the *FMR1 locus* and the trinucleotide repeat size of the resulting fragments was evaluated by electrophoresis in agarose gel.

#### Multiplex ligation-dependent probe amplification (MLPA)

MLPA was conducted to detect abnormal copy-number variations (CNVs) in subtelomeric regions of the chromosomes, as well as frequent interstitial CNVs. MLPA consists on the amplification of different probes using a single PCR primer pair. Each probe detects a specific subtelomeric DNA sequence. Three different kits from MRC-Holland were employed. Two of them (SALSA^®^ MLPA^®^ probemix P036-E3 SUBTELOMERES MIX 1 and SALSA^®^ MLPA^®^ probemix P070-B3 SUBTELOMERES MIX 2B) were used for examining subtelomeric regions. The third one (SALSA^®^ MLPA^®^ probemix P245-B1 Microdeletion Syndromes-1A) was used to detect frequent microdeletion and microduplication syndromes: 1p36 deletion syndrome, 2p16 deletion syndrome, 2q23 deletion syndrome (involving *MBD5*), 2q33 deletion syndrome (involving *SATB2*), 3q29 deletion syndrome, Wolf-Hirschhorn syndrome (resulting from a 4p16 3 deletion), Cri du Chat syndrome (resulting from a 5p15 deletion), Sotos syndrome associated to a 5q35.3 deletion, Williams syndrome (resulting from a 7q11.23 deletion), Langer-Giedion syndrome (resulting from a 8q deletion), 9q22.3 deletion syndrome, 15q24 deletion syndrome, Prader-Willi and Angelman syndromes (resulting from paternal or maternal 15q11.2 deletions), a severe variant of Rubinstein-Taybi syndrome resulting from a deletion in 16p13.3, 17q21 deletion syndrome, Miller-Dieker syndrome (resulting from a 17p deletion), Smith-Magenis syndrome (resulting from a 17p11.2 deletion), NF1 microdeletion syndrome (aka Van Asperen syndrome), linked to a 17q11.2 deletion), Phelan-McDermid syndrome (resulting from a 22q13 deletion), Di George syndrome (resulting from deletions in 22q11, distal 22q11, or 10p14), Rett syndrome (associated to *MECP2* deletion) and *MECP2* duplication syndrome. The PCR products were analyzed by capillary electrophoresis in an automatic sequencer Hitachi 3500 and further analyzed with the Coffalyser V 1.0 software from MRC-Holland.

#### Microarrays for whole-genome CNVs search and chromosome aberrations analysis

The DNA from the patient and her parents was hybridized on a CGH platform (Agilent Technologies). The derivative log ratio spread (DLRS) value was 0.172204. The platform included 60.000 probes. Data were analyzed with Agilent CytoGenomics 3.0.5.1 and qGenViewer, and the ADM-2 algorithm (threshold = 6.0; abs (log2ratio) = 0,25; aberrant regions had more than 4 consecutive probes).

#### Microarrays for DEGs search

In order to determine the genes that could be differentially expressed in the proband (DEGs) compared to her healthy parents, microarray analyses of blood samples from the three of them were performed. Total RNA was extracted with the PAXgene Blood RNA Kit IVD (Cat No./ID: 762164). RNA quality and integrity were confirmed with a Bioanalyzer RNA 6000 Nano. All samples had RNA integrity number (RIN) values above 9. An Affymetrix^®^ Scanner 3000 7G was then used for analyzing transcriptome changes. The resulting raw data were processed with the Affymetrix^®^ GeneChip^®^ Command Console^®^ 2.0 program. Next, *.CEL files were checked to certify the RNA integrity and the suitability of the labeling and the hybridization processes. Finally, the raw data from the different arrays were normalized with the SST (Signal Space Transformation)-RMA (Robust Microarray Analysis) tool (Irizarry et al., 2003). Normalized data (*.CHP files) were subsequently used to search for DEGs in the proband compared to her parents. Statistical analyses were conducted with the LIMMA (Linear Models for Microarray Analysis) package of BioConductor, using the TAC 4.0 software.

## RESULTS

### Clinical History

The proband (Figure 1A) is a girl born by normal delivery after 36 weeks of low-risk gestation. The mother was a 29-year-old woman, who suffered from Crohn’s disease treated with Imurel. At delivery, no signs of disease were observed in the newborn (serology was negative and there was no evidence of group B streptococcus (GBS) infection). Nonetheless, the baby exhibited a supernumerary digit in her right hand, which was subsequently removed by surgery. At birth, her weight was 3.51 kg (percentile 79^th^), her height was 52 cm (percentile 95^th^), and her occipitofrontal circumference was 36 cm (percentile 96^th^). Apgar scores were 9 (at 1’) and 10 (at 5’). When she was 12 months old, her weight was 8,64 kg (percentile 38^th^), her height was 73,5 cm (percentile 42^th^), and her occipitofrontal circumference was 47 cm (percentile 93^th^). At age 1 year and 7 months, her weight was 10 kg (percentile 34^th^), her height was 81 cm (percentile 42^th^), and her occipitofrontal circumference was 48,5 cm (percentile 93^th^).

**Figure 1.**
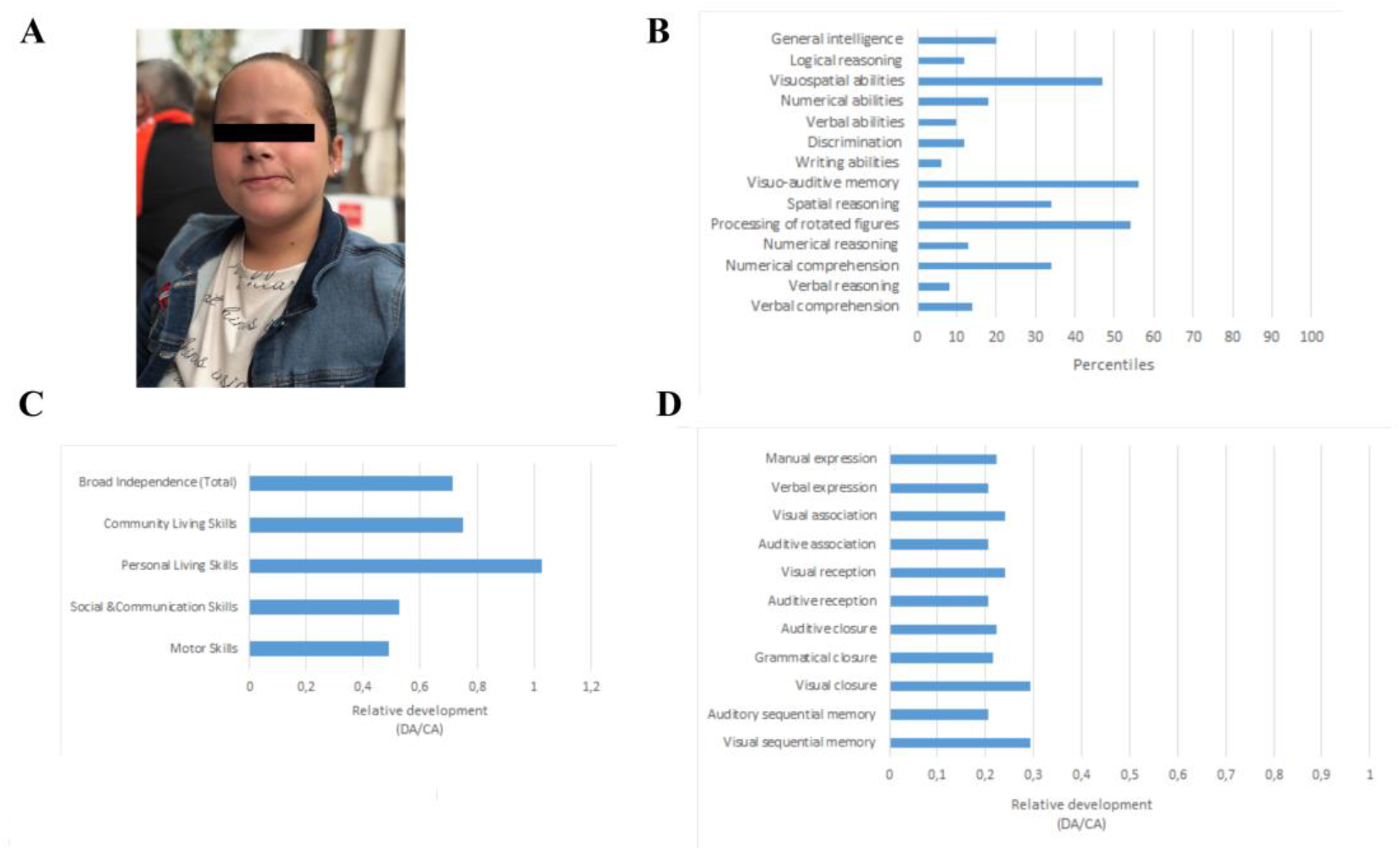
Main physical and cognitive findings in the proband. A. Facial picture of the proband. B. Developmental profile of the proband at the age of 8 years according to the Batería de Aptitudes Diferenciales y Generales (BADyG). C. Developmental profile of the proband at the age of 9 years and 8 months according to the Spanish version of the Inventory for Client and Agency Planning (ICAP). In order to make more reliable comparisons, the resulting scores are shown as relative values referred to the expected scores according to the chronological age of the child. Abbreviations: DA, developmental age; CA, chronological age. D. Developmental profile of the proband at the age of 9 years and 8 months according to the Spanish version of the Illinois Test of Psycholinguistic Abilities (ITPA). Abbreviations: DA, developmental age; CA, chronological age.

The proband’s parents reported on hypotonia that hindered sucking during her first months of life. Hypotonia worsened with time and when she was 12 months old, a cranial magnetic resonance imaging (MRI) was performed. Main findings were a reduced white matter signal-intensity in the temporal lobe of both hemispheres, mostly impacting on the left hemisphere, as well as a mild delay in the myelination pattern of subcortical areas. At the age of 34 months, a prominent forehead (SD <2) was a distinctive feature, as well as 4 hyperchromic spots and abnormally high levels of thyroid-stimulating hormone (TSH). Overall, the girl showed a significant psychomotor delay. At 12 months of age, she had not taken her first steps and showed unstable head control and sitting. Generalized hypotonia, as well as motor and cognitive delay were diagnosed at that age. She was able to stand up and wander with support only at 24 months of age. She also began to manipulate objects at that age. When she was 32 months old, she was able to move from dorsal decubitus to sitting without help. At 34 months of age, she showed typical strength, muscle tone, and spontaneous movements. At this age, she was also able to stand stably and to wander without aid. She succeeded in grabbing objects, passing them from one hand to the other, and in coordinating fine movements. When she was 4 years old, she already walked without aid, but some motor clumsiness and frequent falls were observed. She still suffered from slight hypotonia with kyphosis when sitting without support. When the proband was 6 years old, she was diagnosed with Valgus foot on both feet. She still fell down frequently and reported pain in both soles.

### Early language development

Regarding language development, babbling was first observed at the age of 12 months, whereas bi-syllabic utterances were first reported when the proband was 2 years old. At that age, she already understood simple direct commands. Pointing appeared at the age of 27 months. First words were attested at the age of 32 months. When the proband was 4 years old, she was able to understand complex verbal orders. Nonetheless, expressive language was severely impaired. Accordingly, the two-word stage was reached at the age of 56 months only, with an active vocabulary of around 20 words and the use of non-inflected verbs as a hallmark of her discourse. Telegraphic speech was reported from age 6-7 only.

In spite of her marked language deficits, the proband has being attending a regular school. During the preschool education period, her teachers reported that she mostly communicated through screaming, gesturing, and bringing others to the place where objects of interest were placed, as she was unable to name them. She experienced difficulties with focused attention and she was not interested in new stimuli. She seemed to understand simple commands and sometimes she produced some understandable words. When she was 3 years old she scored significantly low in the Gardner Receptive One Word Picture Vocabulary (ROWPVT), with a verbal age of 2 years and 2 months. At the age of 6, her receptive language was still seriously delayed. She scored 16 direct points in the Peabody Picture Vocabulary Test (PPVT-3) (<1^st^ percentile; −3 SD), which corresponds to a verbal IQ of 55 and a verbal age of 2 years and 7 months. She communicated mostly through facial or body gestures, or touching the other with her hand, as speech and expressive language was seriously impaired. When the proband was 8 years old, her cognitive development was assessed with the Batería de Aptitudes Diferenciales y Generales (BADyG). She scored far below typicality, being the verbal abilities the most impaired domain, whereas spatial abilities were quite preserved (Figure 1B). At the time of our evaluation, when she was 9 years and 8 months old, the proband was attending a normal classroom, although she was assisted by a support teacher and a speech therapist.

### Detailed cognitive and language assessment

At the age of 9 years and 8 months, the proband’s global developmental profile and adaptive behavior were evaluated with the Spanish version of the Inventory for Client and Agency Planning (ICAP). The resulting scores were suggestive of a delay of 3 years and 3 months, being motor skills and communication skills the most impaired domains, whereas personal living skills were the most preserved domain. Community living skills were moderately delayed (Figure 1C; see Supplemental file 1 for details).

Focusing on language, her psycholinguistic development was assessed in detail with the Spanish version of the Illinois Test of Psycholinguistic Abilities (ITPA). The resulting composite psycholinguistic age was 5 years and 2 months, well below her chronological age. Still, she showed a quite irregular profile, with visual and motor abilities being quite spared, but with auditory and grammatical processing abilities, as well as vocal emission, being the most impaired domains (Figure 1D; see Supplemental file 1 for details). More specifically, we found that her receptive language was seriously delayed. She scored 53 direct points in the Peabody Picture Vocabulary Test (PPVT-3), which is suggestive of a verbal age of 5 years and 1 month (see Supplemental file 1 for details). Poor vocabulary is expected to account as well for her problems for understanding language. Likewise, we found that speech production and phonological awareness were severely delayed, exhibiting features that are typically found in children between 4-6 years old, according to the Registro Fonológico Inducido test (see Supplemental file 1 for details). Fluency problems were frequently attested. The proband experienced marked problems for correctly uttering tri-syllabic words, or even bi-syllable words containing consonant clusters. Deletions of weak syllables were frequently observed (e.g. *tuga* for *tortuga* ‘turtle’), as well as cluster reduction (e.g. *peso* for *preso* ‘prisoner’) and assimilations resulting in consonant harmony (e.g. *zaza* instead of *taza* ‘cup’ or *pambor* instead of *tambor* ‘drum’).

Lastly, we examined how the proband put her knowledge of language into use for communicating. We analyzed three different naturally-occurring interactions with her family (see Supplemental file 2 for details). At the expressive level, we confirmed the existence of widespread articulatory problems, including consonant deletions (e.g. *apatos* instead of *zapatos* ‘shoes’; *artera* instead of *cartera* ‘bag’; *e* instead of *él* ‘he’; *Carcola* instead of *Caracola* ‘The Seashell’), deletions of weak syllables (e.g. *tuche* instead of *estuche* ‘case’; *pues* instead of *después* ‘after’; *mano* instead of *hermano* ‘brother’; *jama* instead of *pijama* ‘pyjama’), and simplifications of consonant clusters (e.g. *apoco* instead of *tampoco* ‘neither’; *epués* instead of *después* ‘after’; *pima* instead of *prima* ‘cousin’). We also observed frequent sound assimilations (e.g. *Canme* instead of *Carmen* (proper noun)), non-pervasive frontalizations (e.g. *da* instead of *la* ‘the’; *dlase* instead of *clase* ‘class’), and many instances of loose articulation. Phonetic pharaphasia (e.g. *felfa\** instead of *lazo* ‘bow’; *tunales\** instead of *naturales* ‘natural sciences’) sometimes resulted in idiosyncratic neologisms. Overall, these expressive difficulties seriously hindered the intelligibility of the proband’s discourse. The mean length of utterance in words (MLUw), a common measure of language structural complexity, was 3.9, which is typically found in children aged between 3 years and 6 months and 3 years and 11 months (Rice et al., 2010). Morphosyntactic errors included omissions and substitutions of bound morphemes (e.g. *viene* instead of *vienen*_PL_ ‘they come’; *quién es* instead of *quiénes*_PL_ *son*_PL_ ‘who are they’; *el* instead of *la* ‘the_PEM_’), and omissions of different word classes such as prepositions (e.g. *veintiséis mayo* instead of *veintiséis de mayo* ‘May the 26^th^’; *hablar* instead of *para hablar* ‘to talk’), object pronouns (*pongo* instead of *me pongo* ‘I wear’; *gusta* instead of *te gusta* ‘do you like’), articles (*a piscina* instead of *a la piscina* ‘to the swimming pool’), and auxiliary verbs (*ía* instead of *he ido* ‘I have gone’). Sporadic inversions of the canonical word order in Spanish were also attested (i.e. subject-object-verb instead of subject-verb-object).

In spite of these problems with structural components of language, the proband’s pragmatic skills (i.e. how language is put into use) were significantly spared. First, she mastered turn-taking during conversational exchanges (Sacks et al., 1974), including gaze selection of the interlocutor (even among six participants), or the use of paralinguistic, non-verbal, and echolalic devices for taking the floor. Second, she exhibited signals of metapragmatic awareness, as when feeling embarrassed because of her slow take of the floor during conversational exchanges due to her poor expressive abilities. Third, she seemed to master Grice’s Cooperative Principle (Grice, 1975), including the ability to infer implicit meanings resulting from the flouting of Gricean maxims (Davies, 2007). Accordingly, our proband was able to detect blatant violations of the maxim of quality (‘assume that an utterance will generally be true’) for humorous purposes, and to flout the maxim of quantity (‘give the right amount of information taking into account contextual requirements’) for conveying implicit meanings.

### Cytogenetic and molecular analyses

Routine cytogenetic and molecular analyses of the proband were performed when she was 8 years old. PCR analysis of the FMR1 fragile site resulted normal. MLPAs of selected subtelomeric regions yielded normal results too. Probes for common CNVs failed to detect genomic signatures of high prevalent syndromes resulting from microdeletions or microduplications.

A comparative genomic hybridization array (array-CGH) was then performed. The array-CGH identified a duplication of 1 Mb in the 1q42.3q43 region (arr[hg19] 1q42.3q43 (235,963,632-236,972,276)x3) (figure 2A). The duplicated region includes 16 genes of which 9 are protein-coding genes (figure 2B). The proband also bears a second microduplication in chromosome 1 (arr[hg19] 1q21.2 (148,867,551-149,244,468)x3). The duplicated region contains 6 pseudogenes and 2 genes coding for snRNAs, but no protein-coding genes, and the duplication is predicted as benign.

**Figure 2.**
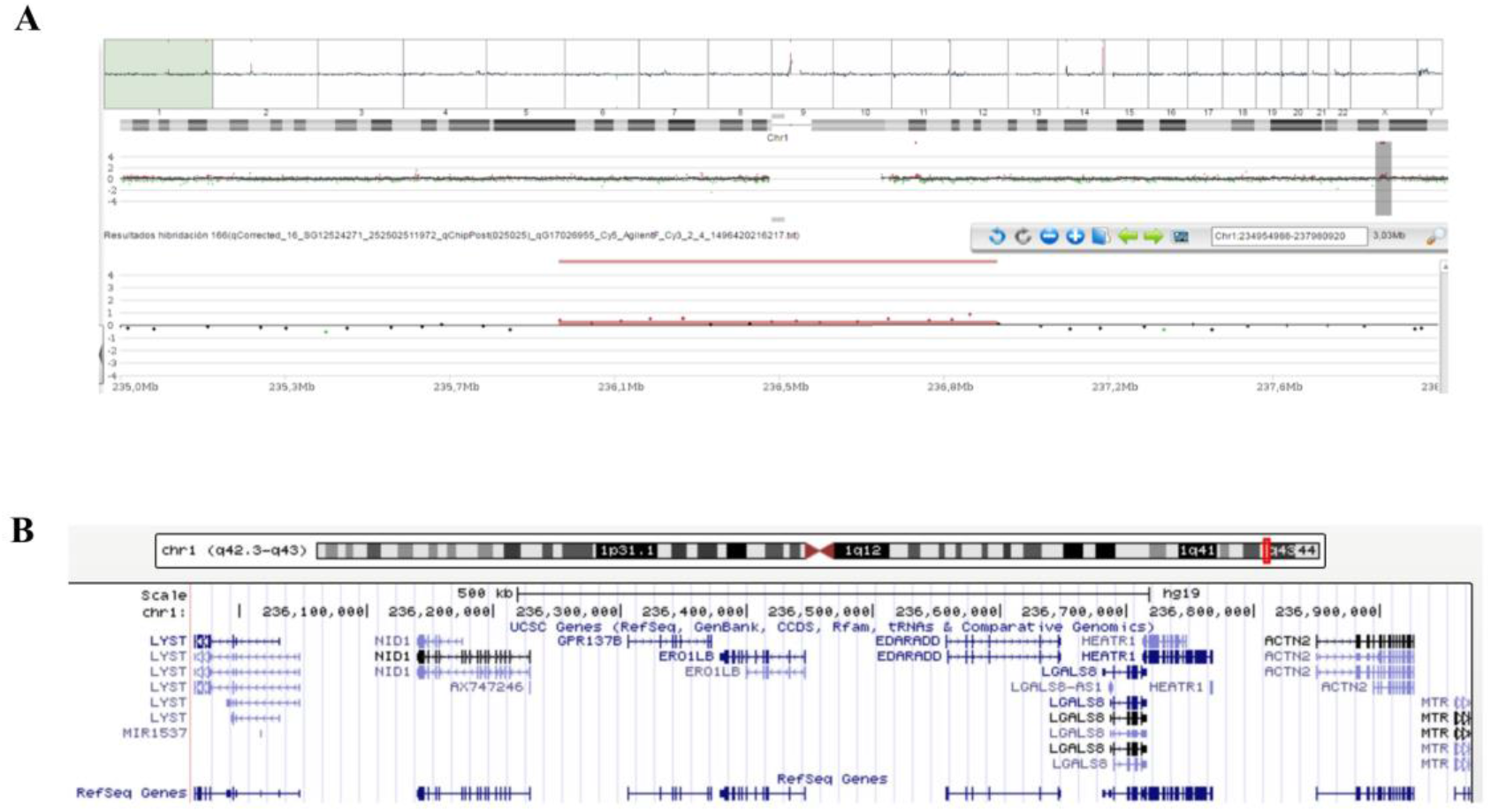
Chromosomal alterations found in our proband. A. Screen capture of the array-CGH of the proband’s chromosome 1 showing the microduplication at 1q42.3q43. B. Screen capture of the UCSC Genome Browser (https://genome.ucsc.edu/) showing the genes duplicated in the proband.

Our proband exhibits some of the physical, behavioral, and cognitive features of partial 1q duplications (Table 1). The duplication in 1q42.3q43 was inherited from the proband’s non-symptomatic father (figure 3A). The man exhibits an average verbal IQ, with no evidence of impairment in any of the assessed domains (figure 3B). Additionally, he exhibits a good command of verbal abilities as assessed with the BAIRES test, around 82^nd^ percentile, but low reading abilities as assessed with the PROLEC-SE test, below 5^th^ percentile, mostly resulting from poor text understanding (figure 3C). He doesn’t show any of the other distinctive features of partial 1q duplications.

**Table 1.**
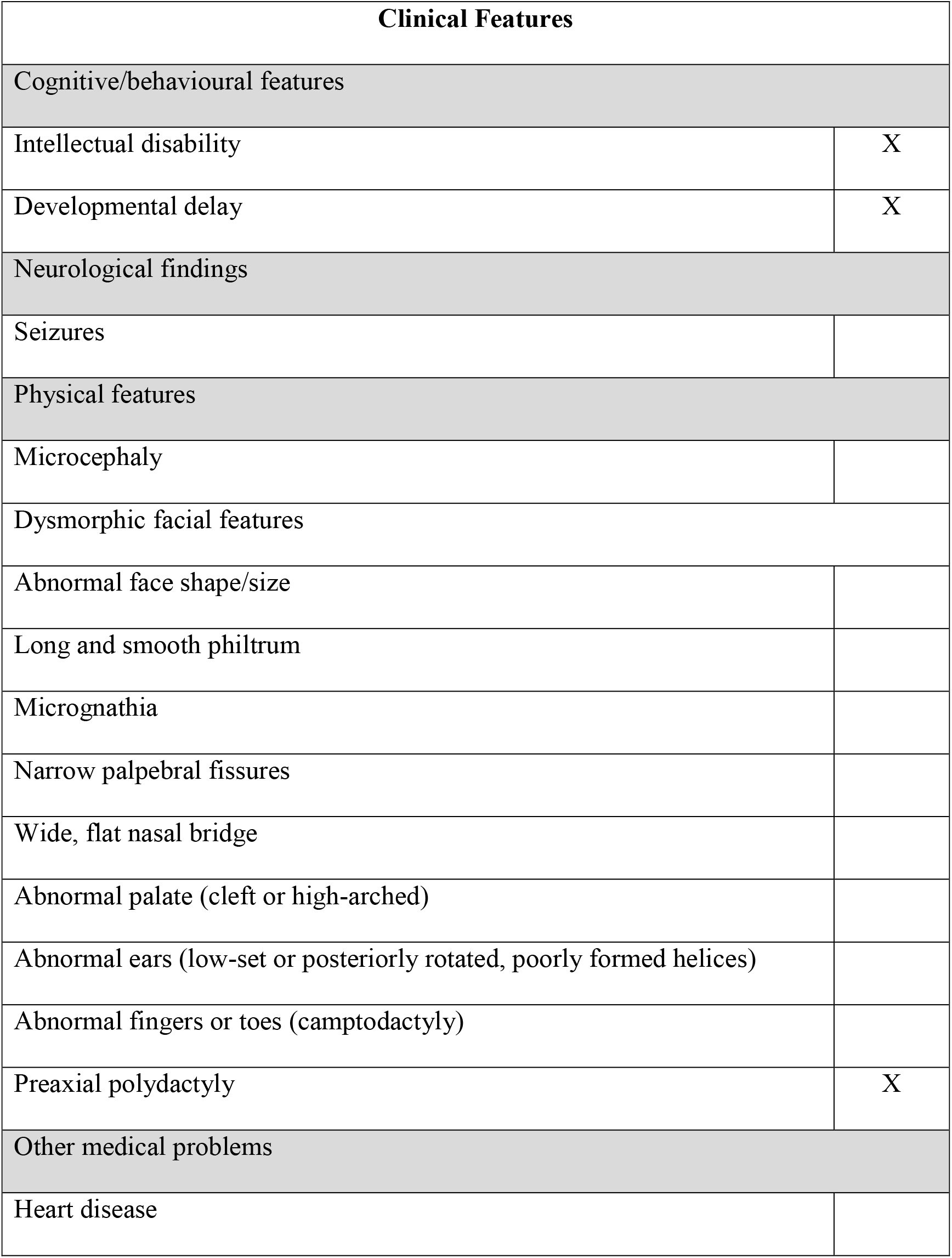
Summary table with the most relevant clinical features of our proband compared to other patients with distal 1q duplications (following Hemming et al. 2016 and Morris et al. 2016)

**Figure 3.**
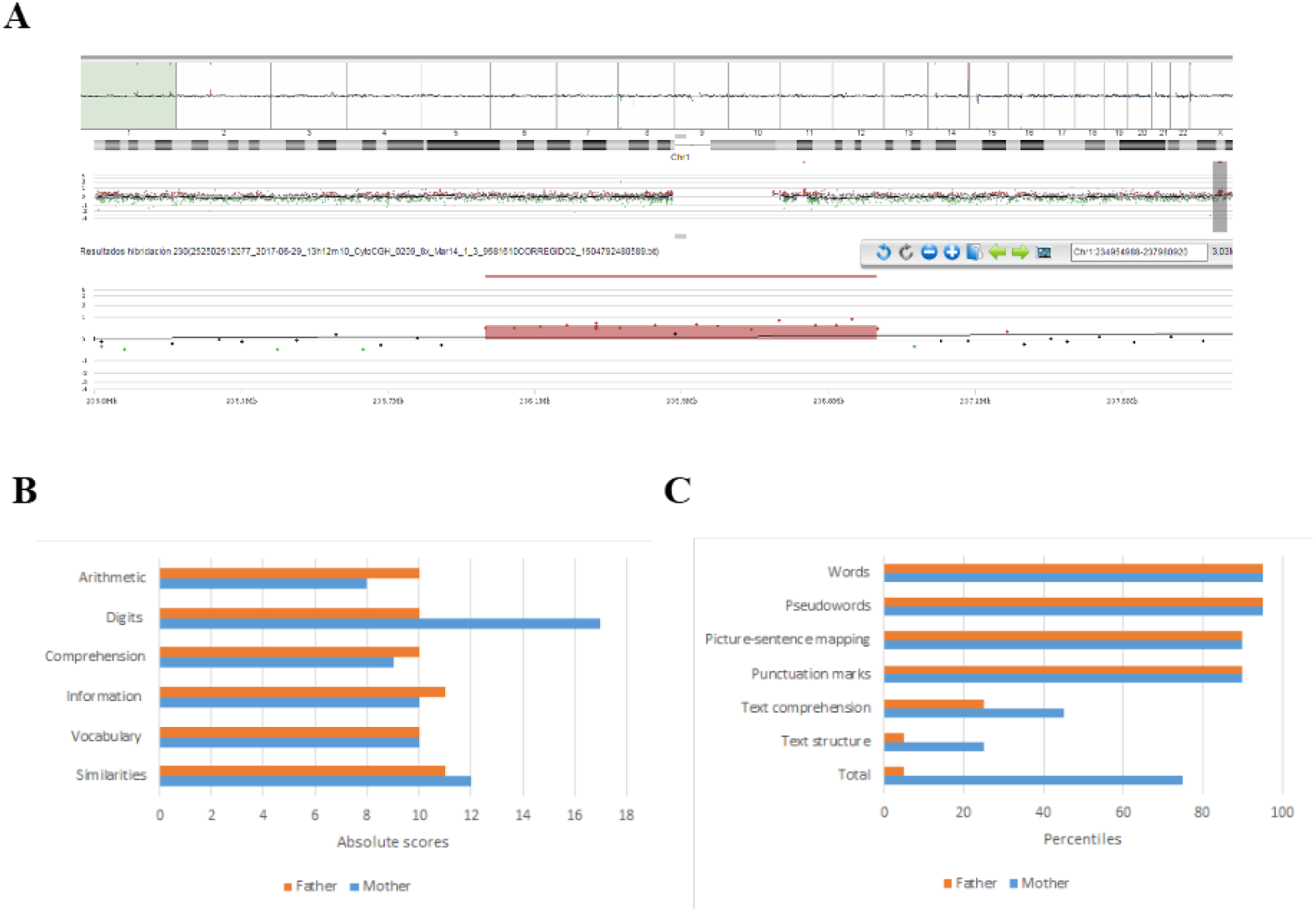
Chromosomal alterations and linguistic profile of the proband’s father. A. Screen capture of the array-CGH of the chromosome 1 of the proband’s father showing the microduplication at 1q42.3q43. B. Developmental profile of the proband’s father according to the verbal component of the WAIS-III (for comparison, the figure includes the profile of the healthy, non-carrier mother). C. Reading abilities of the proband’s father according to the PROLEC-SE test (for comparison, the figure includes the profile of the healthy, non-carrier mother).

In order to delve into the molecular causes of the speech and language problems exhibited by the proband, we conducted several in silico analyses. First, we surveyed the available literature searching for clinical cases linked or associated to the mutation, dysfunction, or dysregulation of the genes duplicated in our proband. We found that a duplication affecting *EDARADD* has been identified in a patient with autism spectrum disorder (ASD) (Prasad et al., 2013). *GPR137B* has been associated to neurocognitive improvement of schizophrenic patients after antipsychotics intake (McClay et al., 2011). *LYST*, which is interrupted by the duplication event, is a candidate for Chediak-Higashi syndrome (OMIM # 214500), a rare autosomal recessive disorder which entails diverse neurologic problems, including cognitive decline and parkinsonism (Kaplan et al., 2008; Introne et al., 2017). Finally, two polymorphism of *MTR*, which is also interrupted by the duplication event, have been associated with bipolar disorder (BD) and schizophrenia (SZ) (Kempisty et al., 2007).

Additionally, we surveyed DECIPHER (https://decipher.sanger.ac.uk/) looking for similar duplication events in other individuals, but also for duplications smaller than the region duplicated in the proband, that might help stablish more robust genotype-to-phenotype links regarding her language disabilities. We found that duplications encompassing *HEATR1, ACTN2*, and *MTR* usually entail delayed motor development, but no language problems (patient 333241). By contrast, duplications encompassing *LGALS8* and affecting *EDARADD* and *HEATR1* normally result in language delay mostly impacting on the expressive domain (patient 248308) (Figure 4).

**Figure 4.**
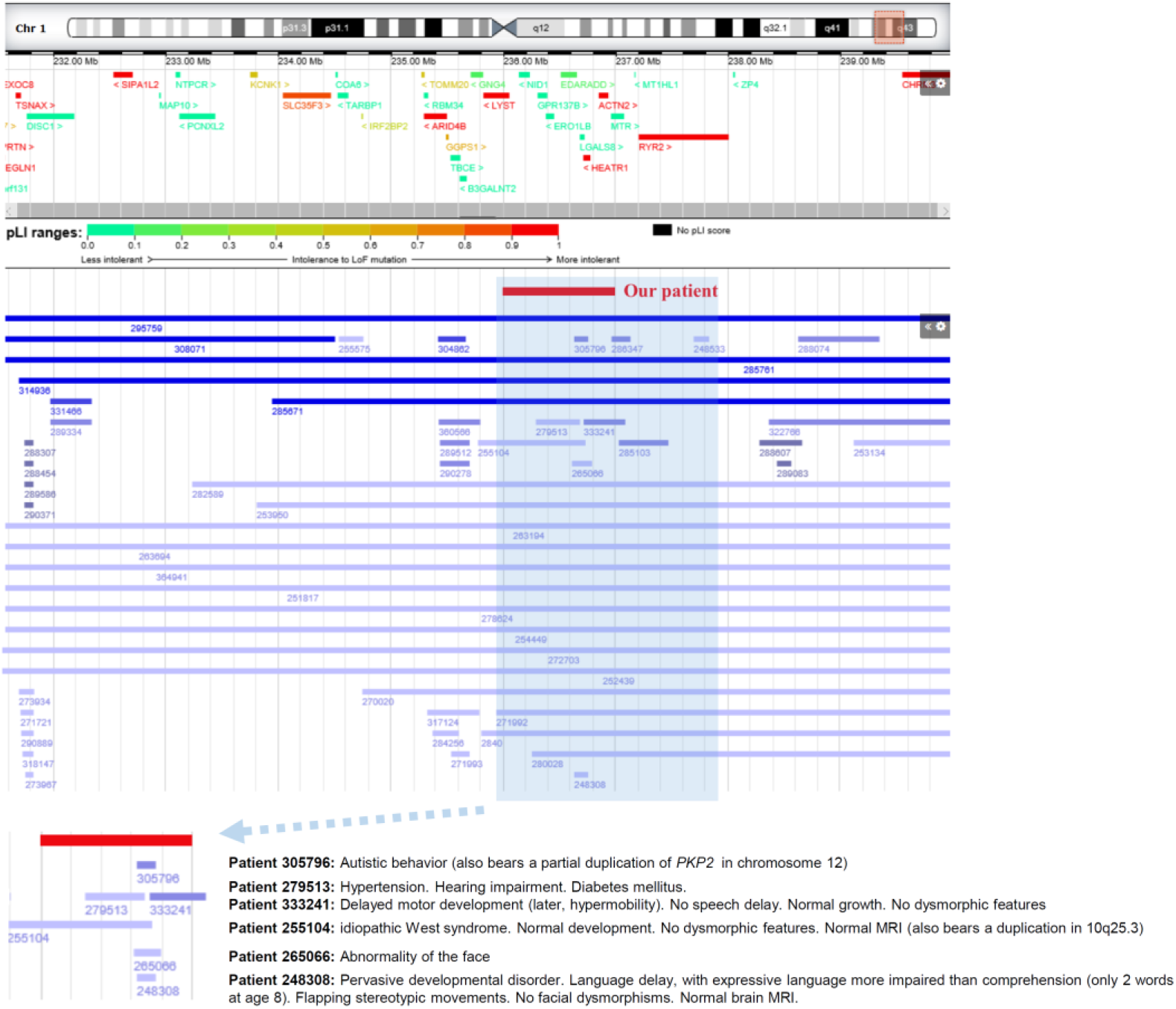
DECIPHER patients of interest bearing chromosomal duplications at 1q42.3q43 with similar or smaller sizes than the one found in the proband. The patients’ phenotype is summarized at the bottom of the figure.

Finally, we used String 10.5 (www.string-db.org) for uncovering potential functional links between the genes duplicated in our proband and genes related to language function, specifically, candidate genes for i) prevalent language disorders (developmental dyslexia (DD) and specific language impairment (SLI), as listed by Paracchini et al. 2016, Pettigrew et al. 2016, and Chen et al. 2017); and ii) language evolution, as discussed by Boeckx and Benítez-Burraco (2014a,b) and Benítez-Burraco and Boeckx (2015) (many of the genes belonging to this second group are also candidates for language dysfunction in broader cognitive disorders, particularly, ASD and SZ, as discussed in detail in Benítez-Burraco and Murphy, 2016, Murphy and Benítez-Burraco, 2016 and Murphy and Benítez-Burraco, 2017). The whole set of genes considered in this analysis is listed in the Supplemental file 3. String 10.5 predicts physical and functional associations between proteins relying on different sources (genomic context, high-throughput experiments, conserved coexpression, and text mining) (Szklarczyk et al. 2015). Several proteins (GRIN2A, GRIN2B, HRAS, PVALB, and ITGB4) were predicted to be direct interactors of the product of one of the genes located within the duplicated fragment, namely, ACTN2 (Figure 5). Co-expression data and experimental/biochemical data further pointed out to a functional link between HEATR1 and NOP9, as well as between LYST and GABARAP.

**Figure 5.**
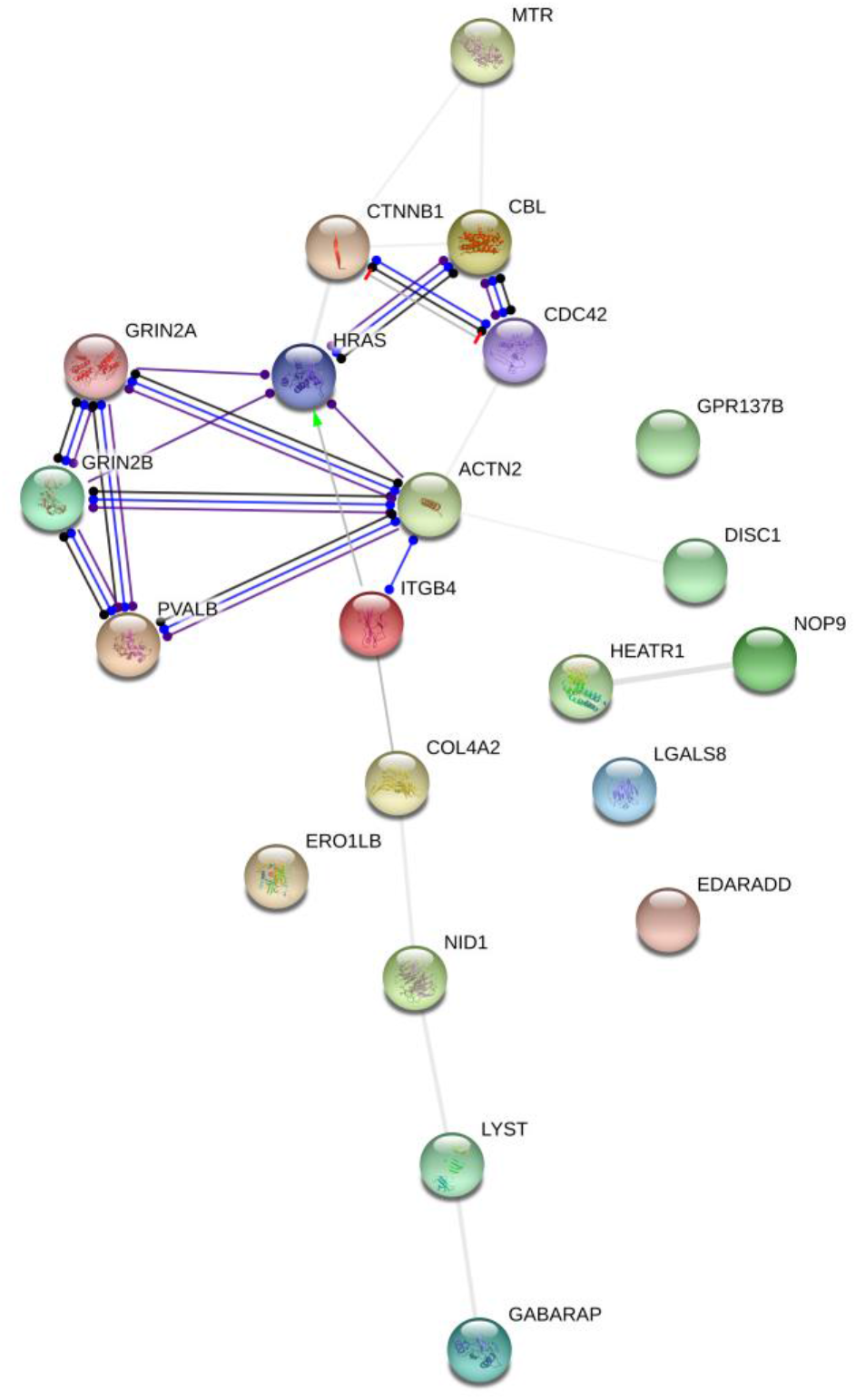
Interaction network of the proteins encoded by the genes duplicated in the proband. The network was drawn with String (version 10.5; Szklarczyk et al. 2015) license-free software (http://string-db.org/), using the molecular action visualization. It includes the products of the protein-coding genes duplicated in the subject, their potential interactors according to String, and the products of 15 strong candidates for language development and/or evolution discussed in the text. Colored nodes symbolize gene/proteins included in the query (small nodes are for proteins with unknown 3D structure, while large nodes are for those with known structures). The color of the edges represents different kind of known protein-protein associations. Green: activation, red: inhibition, dark blue: binding, light blue: phenotype, dark purple: catalysis, light purple: post-translational modification, black: reaction, yellow: transcriptional regulation. Edges ending in an arrow symbolize positive effects, edges ending in a bar symbolize negative effects, whereas edges ending in a circle symbolize unspecified effects. Grey edges symbolize predicted links based on literature search (co-mentioned in PubMed abstracts). Stronger associations between proteins are represented by thicker lines. The medium confidence value was .0400 (a 40% probability that a predicted link exists between two enzymes in the same metabolic map in the KEGG database: http://www.genome.jp/kegg/pathway.html). The diagram only represents the potential connectivity between the involved proteins, which has to be mapped onto particular biochemical networks, signaling pathways, cellular properties, aspects of neuronal function, or cell-types of interest.

Besides these in silico analyses, we also conducted in vitro analyses. Specifically, we performed microarray analyses of blood samples from the proband to determine whether she exhibited altered patterns of gene expression that may account for the observed symptoms. We used her healthy parents as a control. The results of the microarrays are listed in the Supplemental file 4. First, we checked whether the genes highlighted above were differentially expressed in the proband compared to her unaffected parents (Figure 6). Genes showing an opposite expression pattern in the proband and in both healthy parents were of particular interest to us. Among the protein-coding genes located within the duplicated fragment, we found that *EDARADD* is downregulated in our proband, whereas *LYST* is upregulated. Also *ACT2* is slightly upregulated. With regards to the predicted functional partners of the duplicated genes with a known role in language development, language impairment and/or language evolution, we found that neither of them was significantly dysregulated in the girl (i.e. with FC > 2 compared to both healthy parents). Only *HRAS* and *CMIP* could be regarded as slightly upregulated in our subject.

**Figure 6.**
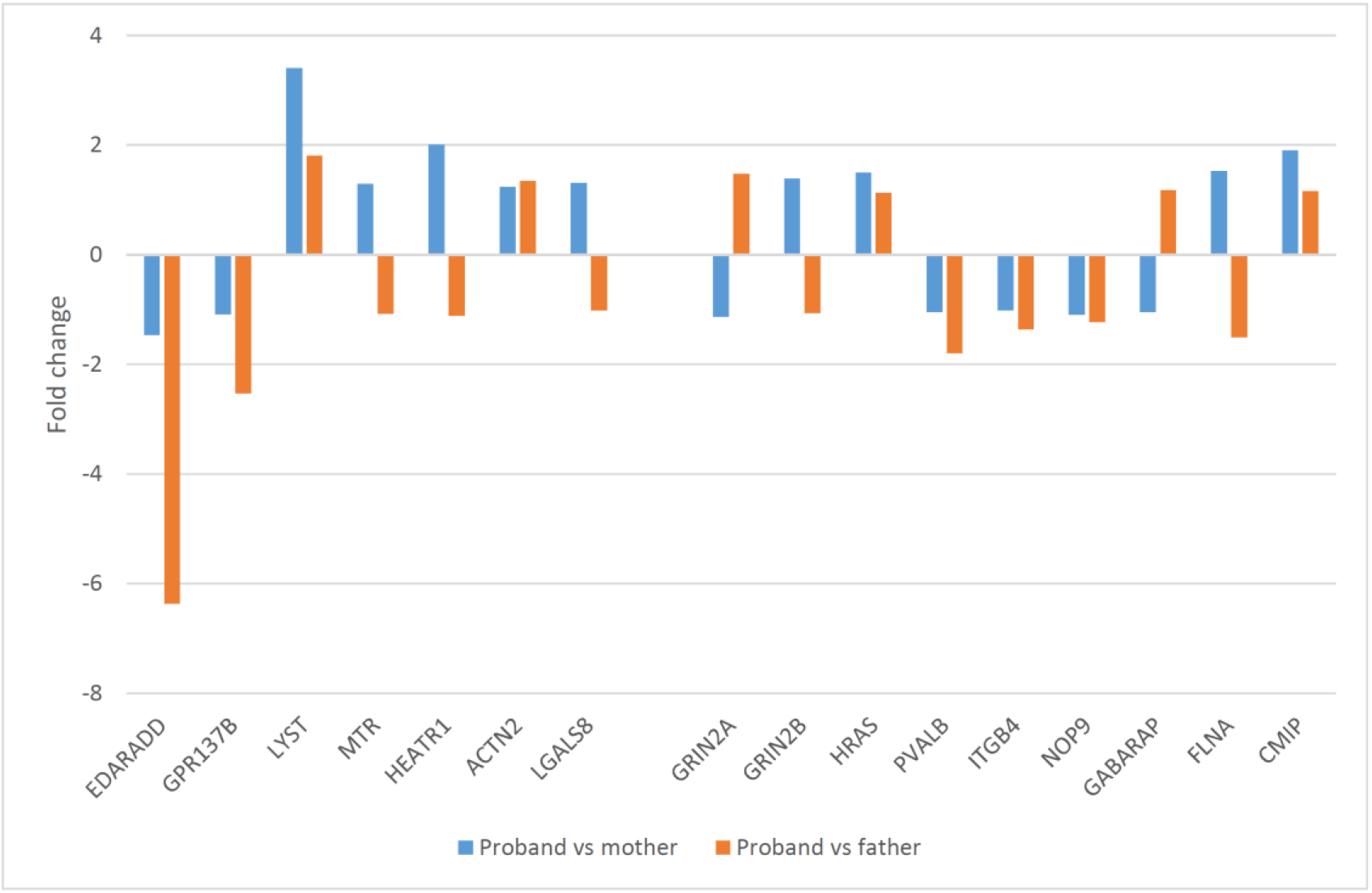
Variation in the expression levels of genes of interest in the proband’s blood compared to her healthy parents (I). The graphic shows the results of the microarray analyses for selected genes within the duplicated fragment and for their predicted functional partners according to String 10.5, as discussed in the text.

Lastly, we searched for additional candidates for the proband’s language deficits, looking for genes exhibiting strong fold changes in our subject (i.e. with FC > 5 compared to both healthy parents). We found several strongly downregulated genes (*IL5RA, MYOM2, DDIT4* and *SLC29A1*), as well as several strongly upregulated genes (*CNTNAP3, WLS*, and *LGALSL*) (Figure 7).

**Figure 7.**
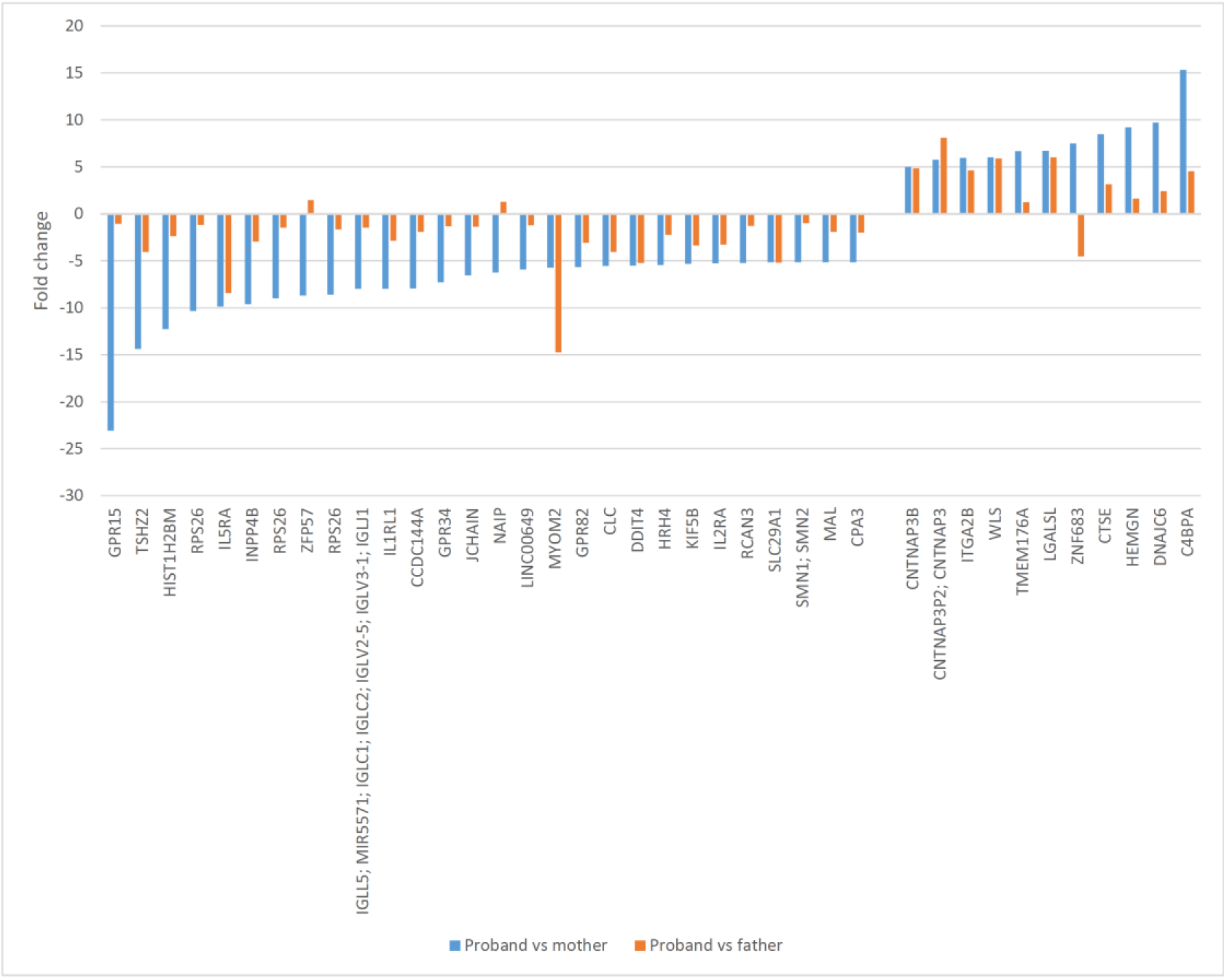
Variation in the expression levels of genes of interest in the proband’s blood compared to her healthy parents (II). The graphic shows the results of the microarray analyses for genes exhibiting fold changes > 5 compared to both healthy parents.

*EDARADD* is expressed at similar levels in all brain areas, whereas *LYST* is expressed at higher levels in the cerebellum, and *ACT2* is mostly expressed in the striatum. Regarding the putative functional partners of the genes located within the duplicated fragment in our proband, *HRAS* is more expressed in the striatum, whereas *CMIP* is similarly expressed in most brain areas. Finally, with regards to the genes that are strongly downregulated in our proband, we found that *MYOM2* and *SLC29A1* are most expressed in the cortex, whereas *DDIT4* is expressed at high levels in the cerebellum and the thalamus, and *IL5RA* is preferentially expressed in the hippocampus. No data were available regarding the three genes that are strongly upregulated in our proband. **Figure 8**. Expression pattern in the blood and the brain of genes of interest. These genes include candidate genes within the duplicated fragment, predicted functional partners, and genes found strongly down- or upregulated in the proband, as discussed in the text. Expression levels of the genes were retrieved from the Genotype-Tissue Expression (GTEx) project (GTEx Consortium, 2013) (https://www.gtexportal.org/home/). Statistical analysis and data interpretation were performed by The GTEx Consortium Analysis Working Group using the Legend: TPM, transcripts per million.

## DISCUSSION

Next generation sequencing facilities, and particularly, comparative genomic hybridization arrays, have increased exponentially the number of genes and chromosomal regions associated to clinical conditions entailing cognitive and language deficits. However, in most cases, no robust genotype-to-phenotype links have been posited. In this paper, we have characterized the cognitive profile of a girl with a microduplication in 1q42.3q43, with a focus on her problems with language. The girl exhibits some of the cognitive, behavioral, and physical features commonly found in people with duplications of the distal 1q (Table 1). She bears one of the smallest duplications of this terminal region of chromosome 1 described to date. Around 20% of DECIPHER patients with a duplication within, overlapping, or encompassing the region duplicated in our proband (N=35) are reported to suffer from language problems. Nonetheless, the duplicated fragments usually encompass more genes than in our proband and/or patients bear other chromosomal rearrangements. Importantly too, assessment of patients usually involves standardized cognitive tests only. Overall, to the best of our knowledge, this is the first detailed report on the language disabilities of a subject bearing only a short distal duplication of the 1 chromosome. For achieving this, we examined the proband’s language strengths and deficits using a battery of specific diagnostic tools, but also conducted fine-grained analyses of language use in natural settings, with a focus on both structural and functional aspects.

In the expressive domain, the proband’s speech is characterized by generalized simplification processes (sound assimilation, consonant cluster reduction, weak syllable deletion), which are typically found in younger children. Because our proband exhibits a noteworthy motor delay, we hypothesize that she suffers from some form of dysarthria. Nonetheless, the results of the RFI test point as well to a phonological impairment, which seemingly contributes to her speech problems. This possibility is reinforced by the low scores obtained by our proband in the ITPA categories for verbal auditory comprehension and association, as well as for sequential auditory memory. These low scores might be pointing to a verbal auditory memory deficit, which is expected to affect the normal acquisition of phonology and ultimately, explain her speech problems, including her tendency to reduce syllabic complexity.

Regarding word morphology and sentence syntax, both are seriously impaired. The MLUw in spontaneous speech is within the range of much younger typically-developing children. Problems are also observed in the receptive domain. Accordingly, her receptive vocabulary is much poorer than expected by age. Since the acquisition of phonology is the scaffold for the acquisition of other components of language, and since comprehension usually goes ahead of expressive abilities during development (Owens, 1984), we contend that the proband’s phonological deficit can be indirectly causing most of her language problems, in both the receptive and the expressive domains.

This remarkable delay in the acquisition of core structural components of language dramatically hinders the child from expressing her needs and feelings, and to a lesser extent, understanding what is happening in her social environment, thus impeding a successful communication with her family and peers. Overall, we hypothesize that this severe language impairment would underlie her learning disabilities and her cognitive deficits. The fact that the proband shows personal living skills that are typical of her chronological age and that enable her to take care of herself supports this view. Moreover, contrary to what has been reported in other developmental disorders resulting in primary intellectual disability, she behaves adequately with strangers, without exhibiting excessive signs of affection. This might also explain the preservation of her pragmatic skills, as she shows metapragmatic awareness and follows the rules governing conversational exchanges, being able to understand and generate conversational implicatures despite her limited expressive resources.

Regarding the molecular causes of the attested problems with language, in silico analyses uncovered several potential candidates for language deficits among the genes located within the duplicated fragment, as well as functional links of interest between some of the genes within the 1q42.3q43 region and robust candidates for language development and/or evolution. Although we have not found evidence of a strong dysregulation of any of these genes, microarray analyses suggest that some of them might be (slightly) up- or downregulated in the blood of our proband. Pretty obviously, pathological changes in

Among the genes located within the duplicated region, one promising candidate is *EDARADD*, which is found significantly downregulated in our proband. This gene encodes an interactor of EDAR, involved in the development of hair, teeth and other ectodermal derivatives. Mutations in *EDARADD* usually result in ectodermal dysplasia (OMIM# 614940; OMIM# 614941) (Headon et al., 2001; Bal et al., 2007; Cluzeau et al., 2019). The gene has been recently associated to a biomarker of ageing, namely, ‘epigenetic age acceleration’, which is predictive of morbidity and mortality (Gibson et al., 2019). It is not known whether the gene plays any significant role at the brain level, where it is expressed at very low levels.

A second promising candidate is *ACTN2*, which is found slightly upregulated in the proband. This gene encodes a bundling protein that anchors actin to different intracellular structures and which contributes to regulate spine morphology and the assembly of the post-synaptic density in neurons (Hodges et al. 2014), as well as the assembly and function of N-methyl-D-aspartate (NMDA) glutamate receptors, particularly in the striatum (Dunah et al., 2000; Bouhamdan et al., 2006). *ACTN2* is predicted to interact with robust candidates for language development, language deficits and/or language evolution, particularly, *GRIN2A, GRIN2B, HRAS, PVALB*, and *ITGB4*. *GRIN2A* and *GRIN2B* encode two components of the subunit NR2 of the NMDA receptor channel, important for long-term potentiation, and ultimately, for memory and learning, and both are candidates for SLI, SZ, and ASD (Murphy and Benítez-Burraco, 2018). Additionally, mutations in *GRIN2A* give rise to epilepsy-aphasia spectrum disorders, including rolandic epilepsies, Landau-Kleffner syndrome, and continuous spike and waves during slow-wave sleep syndrome (CSWSS), which entail speech impairment and language regression (Carvill et al., 2013; Lesca et al., 2013). Speech problems associated to mutations in the gene include imprecise articulation, alteration of pitch and prosody, and dysarthria or dyspraxia (Turner et al., 2015), which recall many of the speech problems observed in our proband. Likewise, mutations in *GRIN2B* have been also found in individuals with cognitive dysfunction and EEG anomalies (Freunscht et al., 2013; Hu et al., 2016). *HRAS* encodes a GTPase important for neural growth and differentiation, long-term potentiation, and synaptic plasticity; the gene is a candidate for Costello syndrome (OMIM# # 218040,) a condition entailing developmental delay and mild to moderate intellectual impairment, with relatively impaired language, particularly in the expressive domain (Axelrad et al., 2011; Schwartz et al. 2013). *PVALB* encodes parvalbumin, a high affinity calcium ion-binding protein with an important role in brain function. Inhibition of parvalbumin-expressing interneurons results in complex behavioral changes, including altered sensorimotor gating, reduced fear extinction, and increased novelty-seeking (Brown et al., 2015). Additionally, the parvalbumin system might represent a convergent downstream endpoint for some forms of ASD, with reduced paravalbumin-expressing neurons resulting in shifting the excitation/inhibition balance towards enhanced inhibition (Filice et al., 2016). Finally, it should be mentioned that inactivating parvalbumin-positive (Pvalb+) interneurons in the auditory cortex alters normal response to sounds (specifically, it strengthens forward suppression and alters its frequency dependence) (Phillips et al., 2017). This is interesting in view of the auditory impairment reported in some patients with similar duplications to the one found in our proband (e.g. DECIPHER patient 279513), but particularly, in view of the verbal auditory memory deficit exhibited by our proband. Lastly, ITGB4 interacts with FLNA (Travis et al, 2004), an actin-binding protein involved in actin cytoskeleton remodeling and neuronal migration (Fox et al., 1998), which in turn binds regarding predictions based on co-expression data and experimental/biochemical data, the links between HEATR1 and NOP9, and particularly, between LYST and GABARAP are also worth considering. *NOP9* is a candidate for language impairment (Pettigrew et al., 2016). *LYST*, which is found slightly upregulated in our proband, has been associated to cognitive decline (Kaplan et al., 2008; Introne et al., 2017). Lastly, *GABARAP* is a candidate for DD and encodes a GABAA receptor-associated protein, which contributes to the clustering of neurotransmitter receptors and to inhibitory neural transmission (Veerappa et al., 2013).

To finish, it is worth considering the biological roles played by the genes that are found more strongly dysregulated in the blood of our proband compared to her healthy parents. Among the genes found strongly downregulated, *IL5RA* is of interest considering the psychomotor problems exhibited by the girl. This gene encodes a subunit of a cytokine receptor and it has been associated to physical activity levels (Letsinger et al., 2019). More interestingly, *DDIT4* encodes a stress-response protein that functions as negative regulator of mTOR, a kinase that regulates cell growth and proliferation, and autophagy, as well as synaptic plasticity. Increased *DDIT4* levels are involved in aspects of neuronal damage, whereas knockdown of *DDIT4* seems to inhibit neuronal apoptosis (Su et al., 2019). More specifically, in rats, overexpression of the gene in the prefrontal cortex results in anxiety- and depressive-like behaviors and neuronal atrophy, whereas mutant mice with a deletion of *DDIT4* are resilient to the behavioral and neuronal consequences of chronic stress (Ota et al., 2014). Abnormally high levels of the gene have been found in the prefrontal cortex of patients with major depressive disorder (MDD) (Ota et al., 2014). The molecular mechanism underlying depressant response in the prefrontal cortex also involves *CACNA1C*. In mice, knockout of *CACNA1C* in this brain region results in antidepressant-like behaviors, whereas overexpression of *DDIT4* reverses this effect (Kabier et al., 2017). *CACNA1C* is a candidate for multiple neuropsychiatric disorders including SZ, BD, and MDD (Ferreira et al., 2008; Green et al., 2010; Curtis et al., 2011; Cross-Disorder Group of the Psychiatric Genomics Consortium, 2013; Kabir et al., 2016). Variants of *CACNA1C* have been associated to deficits in reversal learning (Sykes et al., 2019), whereas some polymorphisms have been associated to decreased semantic verbal fluency in healthy subjects (Krug et al., 2010), as well as to decreased executive function in people with BD (Soeiro-de-Souza et al., 2013). *CACNA1C* expression has been found modulated during associative learning (Sykes et al., 2018). In mice, deletion of *CACNA1C* in glutamatergic neurons gives rise to reduced synaptic plasticity, sociability, and cognition (Dedic et al., 2018). Specifically, haploinsufficiency in the gene results in deficits in prosocial ultrasonic vocalization (Kisko et al., 2018; Redecker et al., 2019). In humans, haploinsufficiency in *CACNA1C* has been related to learning difficulties, expressive language impairment, and motor-skills delay (Mio et al., 2020). By contrast, in humans, *CACNA1C* gain-of-function mutations cause Timothy syndrome (OMIM# 601005), which is featured by syndactyly, autism-like behavior, and language and social delays among other features (Splawski et al., 2004; Napolitano et al., 2015). Several of these features have been attested in our proband. Finally, *SLC29A1* encodes a nucleoside transporter that localizes to the plasma and mitochondrial membranes, and that is involved in both nucleotide synthesis and cytotoxic nucleoside uptake. The gene contributes to basal synaptic transmission, long-term potentiation, neuronal plasticity, and spatial memory (Lee et al., 2018). More specifically, it is involved in glutamatergic neurotransmission (Xu et al., 2015) and the acquisition of goal-directed behavior (Nam et al., 2013). In mice, overexpression of the gene results in changes in dependent behavior, including a greater response to ethanol and a reduced response to caffeine (Kost et al., 2011).

Among the genes found strongly upregulated in our proband compared to her healthy parents, two of them are of particular interest considering her pathological features. *WLS* contributes to regulate the secretion of Wnt signaling molecules, which are involved in different developmental and homeostatic processes (Petko et al., 2019). Considering the motor problems exhibited by our proband, it is of particular interest that WLS is specifically involved in the assembly of the neuromuscular junction between motoneurons and skeletal muscles to control motor behaviour, with mutations in the gene resulting in muscle weakness and neurotransmission impairment (Shen et al., 2018). Interestingly too, *WLS* is also involved in the embryonic development of the cerebellum (Yeung and Goldowitz, 2017). A second promising candidate is *CNTNAP3*, which encodes a type of neurexin involved in cell recognition within the nervous system. This gene is found upregulated in the leukocytes of patients with SZ (Okita et al., 2017). The gene is expressed abundantly in many subcortical regions of the mouse brain, including the striatum, the globus pallidus and the subthalamic nucleus, which are crucially involved in speech and language processing, but also in many other cognitive and emotional functions (Booth et al., 2007; Kotz et al., 2009; Viñas-Guasch and Wu, 2017). In mice, the knockout of *CNTNAP3* results in a delay in motor learning (Hirata et al., 2016), as well as repetitive behaviors, deficits in social interaction, and problems for spatial learning, which recapitulate many ASD-features (Tong et al., 2019). Interestingly too, CNTNAP3 interacts with the synaptic adhesion protein NRLG1, which is also a candidate for ASD and whose mutation results as well in repetitive behaviors and abnormal corticostriatal synapses (Blundell *et al*., 2010).

Finally, we wish to highlight that most of the genes reviewed above are highly expressed in the cerebellum, which is a brain area crucially involved in motor planning, but also in language processing (Vias and Dick, 2017; Mariën and Borgatti, 2018).

## CONCLUSIONS

In conclusion, although the exact molecular causes of language problems (and other cognitive, behavioral and even motor deficits) observed in our proband remain to be fully elucidated, we hypothesize that her distinctive features can result from the altered expression of specific genes within the duplicated 1q42.3q43, with a potential impact on selected functional interactors with a contrasted role in language development and/or evolution. Nonetheless, some other genes outside the duplicated region, that are found strongly dysregulated in the proband, can contribute to the attested impairment in the language domain.

We hope these findings contribute to a better understanding of the behavioral and cognitive phenotype resulting from CNVs of the distal region of the long arm of chromosome 1. A better understanding of the neurobiological foundations of the symptoms found in patients is needed for implementing more efficient pedagogical approaches and intervention strategies aimed to ameliorate their deficits and reinforce their strengths. This is particularly true of low-prevalent conditions, such as this type of CNVs, which are generally understudied and for which good management strategies and intervention tools are not commonly available.

## ACKNOWLEDGMENTS

We would like to thank the proband and her family for their participation in this research. We wish to thank also Dr. Montserrat Barcos Martínez and Dr. Isabel Espejo Portero, from the “Reina Sofía” hospital in Córdoba, for giving us access to the patient and her clinical history. Preparation of this work was supported by funds from the Spanish Ministry of Economy and Competitiveness (grant number FFI2016-78034-C2-2-P [AEI/FEDER,UE] to Antonio Benítez-Burraco, with MFU and SJR as members of the project).

## ETHICS

Ethics approval for this research was granted by the Comité Ético del Hospital “Reina Sofía”. Written informed consent was obtained from the proband’s parents for conducting the psycholinguistic evaluations and the molecular analyses, and for publicizing this case report and all the accompanying tables and images in scientific journals and meetings.

## AUTHORS’ CONTRIBUTION

ABB conceived the paper, performed the molecular experiments, analyzed the genomic/genetic data, and wrote the paper. MFU and SJR conducted the clinical evaluation of the proband, analyzed the psycholinguistic data, and wrote the paper. The three authors approved the final version of the manuscript.

## SUPPLMENTAL DATA

**Supplemental file 1**. Tables with the scores obtained by the proband in the standardized tests used for her assessment.

**Supplemental file 2**. Transcriptions of the conversational exchanges in natural settings used for the analyses of the proband’s language in use.

**Supplemental file 3**. The whole set of genes important for language used for the in silico analysis with String 10.5.

**Supplemental file 4**. Data resulting from the microarray analyses comparing gene expression levels in the blood of the proband and her healthy parents.

**Supplemental file 5**. Developmental expression profiles of genes of interest. These genes include candidate genes within the duplicated fragment, predicted functional partners, and genes found strongly down- or upregulated in the proband, as discussed in the paper. The expression data are from the Human Brain Transcriptome Database (http://hbatlas.org/). Six different brain regions are considered: the cerebellar cortex (CBC), the mediodorsal nucleus of the thalamus (MD), the striatum (STR), the amygdala (AMY), the hippocampus (HIP) and 11 areas of neocortex (NCX).

