## Supplemental file 2 for "Language impairment with a microduplication in 1q42.3q43"

**CONVERSATIONAL ECHANGE 1**

@Begin

@Languages:	spa

@Participants:	CHI Informant, FAT Father, MOT Mother

@Options:	CA

@Transcriber:	Maite Fernández Urquiza

@Time Duration:	00:01:44

@Date:	DEC-2017

@Location:	Andújar

@Situation:	Mealtime at home

@Activities:	CHI gives a gift card to her father

*FAT:	ábrela .

*CHI:	va:le@i [=! starts opening the envelope] <tuya es

[=! deleted /s/ + open vowel sound, dialectal]> [=! shouting] .

%xepr:	$i5:PIM:PMQT:ADQ

*FAT:	vale@i pero léemela !

*CHI:	va:le@i .

*FAT:	<a ver> [?] (.) l(a) has [=! deleted /s/, dialectal] hecho tú?

*CHI:	no [=! showing the card to FAT] [=! shakes head no] .

%xepr:	$et3:GSA

*FAT:	qué pone ahí ?

*CHI:	tú (.) <una (.) [=! pointing to a picture on the card] rasca [/]

rasca> [*] [=! aspirated /s/, dialectal] [=! scratch illustrator] !

%err:	una rasca=rasca una $SYN

%xepr:	$i5:MMN:ORD:INV; $i5:MMN:REP; $et3:GSA

*FAT:	a ver@i [=! laughs] que rasque [=! aspirated /s/, dialectal] una

[=! looking closely to the card] ?

*CHI:	sí (.) hoy .

*MOT:	solo una ?

*CHI:	sí (.) <mañana [=! syllabified] otra> [/] mañana otra

[=! regulator of rythm with hand] .

%xepr:	$i5:MMN:REP; $et3:GSA

*FAT:	a ver@i [=! scratching the card with a fork] .

*CHI:	a xxx !

*FAT:	<euro yo no tengo ni uno> [?] .

*CHI:	no [>] .

*FAT:	no [<] no se rasca [=! aspirated /s/, dialectal].

*CHI:	que sí: !

%sit:	FAT finishes scratching the card and raises it in order to read it.

*CHI:	gusta [*] [=! aspirated /s/, dialectal] o no ?

%err:	gusta=te gusta $MOR $SYN

%xepr:	$i5:MMN:ORD:OMI

*FAT:	a ver qué pone +"/.

*FAT:	+" vale: (.) por una: (.) peli juntos „ sí ?

*CHI:	sí (.) <a ver otra> [>] .

*FAT:	<qué más [=! deleted /s/, dialectal] dice ahí> [?] [<] ?

*FAT:	+" vale por cien (.) mil besos

[=! deleted /s/ in final position, dialectal] .

*FAT:	+" vale por tu cena favorita „ sí ?

*CHI:	0 [=! smiles and nods]. [+ trn]

%xepr:	$et3:GSA

*FAT:	+" vale por tu súper abrazo [=! looks at CHI].

*CHI:	esta [=! aspirated /s/, dialectal][=! points to the card smiling].

*FAT:	+" vale po:r un desayuno en la cama [=! laughs] .

*FAT:	ahí no llego ya (.) lo demás sí (.) el cine vale (.) diez mil besos

[=! deleted /s/ in final position, dialectal] (.) o más

[=! deleted /s/, dialectal] (.) y vale por tu cena favorita también .

*CHI:	y ese [=! points to the card and looks to FAT in the eyes] ?

*FAT:	y ese: +"/.

*FAT:	+" vale por un súper abrazo .

*FAT:	dame un abrazo [=! hugs and kisses CHI].

*CHI:	gusta [=! aspirated /s/, dialectal] ?

%err:	gusta=te gusta $MOR $SYN

%xepr:	$i5:MMN:ORD:OMI

*FAT:	<cuántos besos> [=! deleted /s/ in final position, dialectal] son

(.) diez mil ?

*FAT:	un <dos tre:s> [=! deleted /s/, dialectal][=! kissing CHI ] +//

&*CHI: 0 [=! laughs] .

*CHI:	gusta [=! aspirated /s/, dialectal] o no ?

%err:	gusta=te gusta $MOR $SYN

%xepr:	$i5:MMN:ORD:OMI

*FAT:	sí está [=! aspirated /s/, dialectal] mu@d bonito .

@End

**CONVERSATIONAL ECHANGE 2**

@Begin

@Languages:	spa

@Participants:	CHI Informant, FAT Father, MOT Mother, GFA Grandfather, GMA

Grandmother, BOY Familiar

@Options:	CA

@ID:	spa|CHROMOLANG|CHI|||||Informant|||

@ID:	spa|CHROMOLANG|FAT|||||Father|||

@ID:	spa|CHROMOLANG|MOT|||||Mother|||

@Transcriber:	Maite Fernández Urquiza

@Time Duration:	00:06:22

@Date:	13-MAY-2018

@Location:	Andújar

@Situation:	Family meeting in a restaurant's terrace

@Activities:	Conversation about CHI's first communion party

*CHI:	<veintiséis [=! deleted /s/, dialectal]

mayo> [*] (.) <pongo el> [>] +/.

%err:	veintiséis mayo=veintiséis de mayo $MOR $SYN

%xepr:	$i5:MMN:ORD:OMI

*MOT:	qué haces [=! deleted /s/, dialectal] ?

*CHI:	pongo [*] e:h el vestido [=! aspirated /s/, dialectal] (.) <em apato>

[*] (0.3) viene [*] todos [=! deleted /s/, dialectal] .

%err:	pongo=me pongo $MOR; em apato=los zapatos $PHO $MOR; viene=vienen

$MOR $SYN

%xepr:	$i5:MMN:ORD:OMI; $et3:DRA:FIL; $i5:MMN:ORD:SST; $i5:MMN:ORD:SST

*MOT:	quién te va a peinar [=! deleted /r/, dialectal] ?

*CHI:	ah@i Canme [*].

%err:	Canme=Carmen $PHO

*MOT:	sí (.) y la cabeza: qué vas [=! deleted /s/, dialectal] a llevar

[=! deleted /r/, dialectal] ?

*CHI:	&ar (0.3) +/.

%xepr:	$et3:DRA:DIS

*GMA:	una corona .

*CHI:	&za &fa felfa [*p:n] [?] ?

%err:	felfa=lazo $PHO $LEX

%xepr:	$et3:DRA:DIS; $et3:DRA:DIS

*GFA:	no: corona .

*MOT:	no: dile un lazo .

*CHI:	xxx [=! hands over her mouth].

*GMA:	pero dilo fuerte que no se oye .

*CHI:	lazo [=! syllabified] .

*MOT:	y qué más [=! deleted /s/, dialectal] ?

*CHI:	viene [*] todos [=! deleted /s/, dialectal] .

%err:	viene=vienen $MOR $SYN

%xepr:	$i5:MMN:ORD:SST

*GMA:	quiénes [=! deleted /s/, dialectal] vienen ?

*CHI:	www

%sit:	CHI says a list of nouns helped by her family

*MOT:	y dónde vamos [=! deleted /s/, dialectal] a comer

[=! deleted /r/, dialectal] ?

*CHI:	aquí fuera [=! tapping the table with her hand] .

*MOT:	aquí fuera y <esto dónde estamos>

[=! aspirated /s/ in postnuclear position; deleted /s/ in final position

of the word, both dialectal] ?

*CHI:	hum@i [=! looking around] .

%xepr:	$i5:MQT:RVP:INT

*GFA:	Caracola Caracola Caracola [=! whispers trying to help CHI].

*GMA:	cállate [=! to GFA] .

*FAT:	Caramón [=! hand over his mouth, pretending to speak only to CHI] .

*CHI:	Caramón .

%xepr:	$i5:MQL:ASW:ECO

*MOT:	Caramón [=! laughs] ?

*GMA:	en el camión [=! laughs] ?

*CHI:	camión [=! whispered, laughing out loud] .

%xepr:	$i5:PIM:PMQL:ACOM:ECO

*BOY:	a ver@i Ana que te cuento un secreto (.) pero no se lo digas

[=! deleted /s/, dialectal] a nadie .

*CHI:	0 [=! leans forward, looks to BOY]. [+ trn]

%xepr:	$et3:GSA

*BOY:	sabes [=! deleted /s/, dialectal] quién va a hacer

[=! deleted /r/, dialectal] la comunión contigo ?

*CHI:	quién ?

*BOY:	xxx [>]

*GFA:	la Caracola [<].

*CHI:	0 [=! leans back laughing out loud]. [+ trn]

*BOY:	le dejo un traje mío de la comunión <xxx xxx xxx> [>].

*CHI:	que no: [=! shouting, showing EBM,looks to MOT] !

%xepr:	$et3:GSA

*MOT:	(es)cucha (.) entonses [=! deleted /s/ in final position, dialectal]

qué ?

%sit:	there is people around and CHI gets distracted

*MOT:	Ana (.) entonses [=! deleted /s/ in final position, dialectal] qué

vamos [/-] [=! deleted /s/, dialectal] dónde (.) al Ramón ?

*CHI:	sí al Ramón (.) y epués [*] [=! deleted /s/, dialectal] vamos

[=! deleted /s/, dialectal] a misa .

%err:	epué=después $PHO

*CHI:	y epués [*] [=! deleted /s/, dialectal] vamos [*s:r]

[=! deleted /s/, dialectal] aquí .

%err:	epué=después $PHO vamo=venimos $LEX

%xepr:	$i5:MMN:ORD:SST

*BOY:	a comer [=! deleted /r/, dialectal] .

*CHI:	a comer [=! deleted /r/, dialectal] .

%xepr:	$i5:MQT:RUT:ECO

*BOY:	qué vamos [=! deleted /s/, dialectal] a comer

[=! deleted /r/, dialectal] ?

*CHI:	y epués [*] [=! deleted /s/, dialectal] vamos

[=! deleted /s/, dialectal] a jugar [=! deleted /r/, dialectal] .

%err:	epué=después $PHO

*CHI:	<vamos [/] vamos> [=! deleted /s/, dialectal] <a: piscina> [*] (.) a

bañar [*] [=! deleted /r/, dialectal] .

%err:	a piscina=a la piscina $MOR bañá=bañarnos $MOR $SYN

%sit:	there is a bee and FAT tells boy to kill it.

%xepr:	$i5:MMN:REP; $i5:MMN:ORD:OMI; $i5:MMN:ORD:OMI

*FAT:	www.

*MOT:	www.

*GFA:	www.

*MOT:	venga@i qué más [=! deleted /s/, dialectal] „ Ana ?

*CHI:	e:h +/.

%xepr:	$et3:DRA:FIL

*GMA:	vamos [=! deleted /s/, dialectal] a echar

[=! deleted /r /, dialectal] d(e) eso cómo se llama ?

*CHI:	qué ?

*GMA:	eso que sube p(ara)@d arriba .

*BOY:	<fuegos artificiales> [=! deleted /s/, dialectal] .

*MOT:	y va [/-] qué va a haber [=! deleted /r/, dialectal] ?

*MOT:	va a haber [=! deleted /r/, dialectal] chuches

[=! deleted /s/, dialectal] ?

*CHI:	no sé [=! shrugs] .

*MOT:	no sabes [=! deleted /s/, dialectal] qué te van a regalar

[=! deleted /r/, dialectal] ni na(da)@d ?

*CHI:	sí .

%xepr:	$sa2:ISA:ICOM

*MOT:	a ver@i [=! deleted /r/, dialectal] qué te van a regalar

[=! deleted /r/, dialectal] ?

*CHI:	e:h <tú u:n> [>] [=! pointing to FAT] +/.

%xepr:	$et3:DRA:FIL

*GMA:	una Play+Cuatro [<].

*CHI:	no: [=! looking at GMA, showing EBM] .

%xepr:	$et3:GSA

*FAT:	no le gusta [=! aspirated /s/, dialectal].

*BOY:	no ?

*CHI:	no [=! to BOY, shakes head no] .

%xepr:	$et3:GSA

*MOT:	a ti te gusta [=! aspirated /s/, dialectal] la Play+Cuatro ?

*CHI:	no [=! shakes head no] !

%xepr:	$et3:GSA

*MOT:	qué te gusta [=! aspirated /s/, dialectal] a ti?

*FAT:	la Play+Cinco [=! joking] .

*CHI:	no (.) apoco [*] [=! turns to FAT, taps him in the arm, smiles] .

%err:	apoco=tampoco $PHO

%xepr:	$i5:PIM:PMQL:ACOM

*GMA:	<di que no> [?] .

*GFA:	la ocho [=! joking] !

*CHI:	apoco [*] !

%err:	apoco=tampoco $PHO

*MOT:	qué te gusta [=! aspirated /s/, dialectal] a ti?

*FAT:	un patinete elétrico@d .

*CHI:	sí [=! turns to FAT, points to him, nods, laughs] .

%xepr:	$et3:GSA

*GFA:	www.

*FAT:	www.

%sit:	they speak simultaneously and unintelligibly

*BOY:	un móvil [=! deleted /l/, dialectal].

*GMA:	no: !

*CHI:	sí un móvil [=! deleted /l/, dialectal]

[=! raises her forefinger to attract FAT's attention, laughs] !

*MOT:	pa(ra)@d qué quieres [=! deleted /s/, dialectal] un móvil

[=! deleted /l/, dialectal] ?

*CHI:	hablar [*] [=! deleted /r/, dialectal] on [*] <mis amigos>

[=! deleted /s/, dialectal] .

%err:	hablá=para hablar $MOR $SYN on=con $PHO

*MOT:	q(ué) amigos [=! deleted /s/, dialectal] ?

*BOY:	algo más [=! deleted /s/, dialectal] grande (.) quieres

[=! deleted /s/, dialectal] una tablet [=! deleted /t/, dialectal]

nueva ?

*CHI:	no [!] [=! leans back, arms up towards BOY] ya tengo .

*GMA:	venga@i <qué quier> [/-] qué <quieres más>

[=! deleted /s/, dialectal] de regalo ?

*CHI:	e:h un tuche: [*] +/.

%err:	tuche=estuche $PHO

%xepr:	$et3:DRA:FIL

*FAT:	un pony [=! joking] .

*CHI:	no un pony no [=! shouting] !

*BOY:	xxx xxx xxx xxx xxx .

*CHI:	no (.) a:h una: [/-] un ga:to .

%xepr:	$et3:DRA:FIL; $i5:MMN:REF; $i5:PIM:PMQL:ADQ

*MOT:	un gato [/] un gato ?

*CHI:	sí [=! nods, smiles] .

%xepr:	$et3:GSA

*GMA:	<lo que nos faltaba ya también> [?] .

*CHI:	es [=! deleted /s/, dialectal] mentira es

[=! deleted /s/, dialectal] mentira [=! showing EBM] .

%xepr:	$et3:GSA

*MOT:	ah@i de mentira .

*CHI:	xxx xxx .

*GFA:	no pero un pony sí .

*CHI:	no pony no !

*GFA:	pa(ra)@d montarte .

*MOT:	www.

*WOM:	www.

*BOY:	www.

%sit:	they speak simultaneously and unintelligibly

*CHI:	www.

%sit:	starts telling names of guests again.

*GFA:	y <los otros abuelos> [=! deleted /s/, dialectal] ?

*CHI:	<<quién es> [*] los otros abuelos> [=! deleted /s/, dialectal]

[=! showing EBM] ?

%err:	quién es=quiénes son $MOR $SYN

%xepr:	$i5:MMN:ORD:SST; $i5:MMN:ORD:SST; $et3:GSA

*BOY:	ya los [=! deleted /s/, dialectal] ha dicho .

*CHI:	ya: [=! nods] <los [=! deleted /s/, dialectal] he dicho> [>] .

%xepr:	$et3:GSA;

*MOT:	<cuántos abuelos tienes> [=! deleted /s/, dialectal] ?

*CHI:	0 [=! three EBM] . [+ trn]

%xepr:	$et3:GSA;

*MOT:	tres [=! deleted /s/, dialectal] ?

*GFA:	no: .

*BOY:	xxx xxx xxx [>] .

*CHI:	ella [=! pointing to GMA] e [*] [=! pointing to GFA] [<] cuatro

[=! four EBM] .

%err:	e=él $PHO

%xepr:	$et3:GSA;

*GMA:	uy@i !

%sit:	a man is about falling down

*MOT:	cuántos [=! deleted /s/, dialectal] [>] ?

*CHI:	cuatro [<] [=! four EBM] .

%xepr:	$et3:GSA;

*MOT:	cómo se llaman ?

*CHI:	a [*] Mari e [*] Pepe [=! sighs, counts with fingers] e [*] abuela

Antonia [=! points to GMA] e:l [=! points to GFA, laughs] +//.

%err:	a=la $PHO; e=el $PHO; e=la $PHO

%sit:	everybody laughs

*BOY:	Ana ‡ qué quieres [=! deleted /s/, dialectal] que te regale ?

*GFA:	se quiere <quedar conmigo> [=! laughs] [>].

*CHI:	l(o) que tú quieras [=! deleted /s/, dialectal]

[=! raises her arm toward BOY] .

*BOY:	lo que yo quiera no: lo que xxx xxx .

*CHI:	l(o) que tú quieras [=! deleted /s/, dialectal]

[=! claps once, showing EBM toward BOY] !

%xepr:	$et3:GSA

*CHI:	e:h a ver@i +/.

%xepr:	$et3:DRA:FIL

*MOT:	y dónde vas [=! deleted /s/, dialectal] a dormir

[=! deleted /r/, dialectal] ?

*CHI:	ah@i [=! points to BOY] .

%xepr:	$i5:MQT:RVP:INT

*BOY:	en la cama de arriba [?] con la abuela [=! joking] .

*GMA:	ea@i conmigo !

*CHI:	0 [=! keeps pointing to BOY]. [+ trn]

%xepr:	$et3:GSA

*BOY:	<conmigo no> [!] yo duermo en una cama aparte .

*CHI:	no (.) una aparte no

[=! gesture of disagreement with hand, leans back] !

%xepr:	$et3:GSA

@End

**CONVERSATIONAL ECHANGE 3**

@Begin

@Languages:	spa

@Participants:	CHI Informant, MOT Mother, BOY Brother, FAT Father

@Options:	CA

@ID:	spa|CHROMOLANG|CHI|||||Informant|||

@ID:	spa|CHROMOLANG|MOT|||||Mother|||

@ID:	spa|CHROMOLANG|BOY|||||Brother|||

@Transcriber:	Maite Fernández Urquiza

@Time Duration:	00:03:43

@Date:	05-JUN-2018

@Location:	Andújar

@Situation:	Conversation at home

*MOT:	que le cuente [=! deleted /s/, dialectal] lo que hiciste

[=! aspirated /s/, dialectal] ayé [=! deleted /r/, dialectal] nel

colegio .

*CHI:	ah@i he hecho xxx xxx xxx xxx .

*CHI:	y: [/] y epués [*] [=! deleted /s/, dialectal] (..) he hecho

eligión (.) <pués [*] mates>

[=! deleted /s/ at the end of both words, dialectal] (.) pués [*]

[=! deleted /s/, dialectal] patio (...) y: (0.6) tunales [*]

[=! deleted /s/, dialectal].

%err:	epués=después $PHO; pués=después $PHO; pués=después $PHO;

tunales=naturales $PHO $LEX

%xepr:	$i5:MMN:REP;

*CHI:	y: (..) no sé lo otro (.) no lo sé.

%xepr:	$et3:DRA:ERQ

*MOT:	has [=! deleted /s/, dialectal] dicho: ?

*CHI:	tunales [*] [=! syllabified, deleted /s/, dialectal] .

%err:	tunales=naturales $PHO $LEX

*MOT:	natura:les [=! deleted /s/, dialectal].

*CHI:	no sé: xxx xxx xxx lo otro .

%xepr:	$et3:DRA:ERQ

*MOT:	cuál es [=! deleted /s/, dialectal] lo otro ?

*MOT:	yo no sé cuál es [=! deleted /s/, dialectal] lo otro.

*CHI:	ni yo [=! shakes head no] (0.6).

%xepr:	$et3:GSA

*MOT:	venga@i cuéntale <más cosas>

[=! deleted /s/ at the end of both words, dialectal] que el día <es

[=! deleted /s/, dialectal] muy> [/-] fue muy largo !

*CHI:	<y &e> [/-] y a da [*] tarde (..) ía [*] a dlase [*] y epués [*]

[=! deleted /s/, dialectal] +/.

%err:	da=la $PHO; ía=he ido $PHO; dlase=clase $PHO; epués=después $PHO

%xepr:	$i5:MMN:REF;

*MOT:	cuéntale cómo se llamaba (.) la: profesora de clase .

*CHI:	Mari+Jose .

*MOT:	y cómo es [=! deleted /s/, dialectal] ?

*CHI:	apa [*] .

%err:	apa=guapa $PHO

*MOT:	venga@i (.) descríbesela [=! aspirated /s/, dialectal](.) porque xxx

no la conoce (0.6) venga@i !

*CHI:	xxx e:s [=! deleted /s/, dialectal] (...) no m'acuerdo (0.7).

%xepr:	$et3:DRA:ERQ

*MOT:	bueno cuéntale qué es [=! deleted /s/, dialectal] lo que hiciste

[=! apirated /s/, dialectal] en la comunión a ver@i !

*CHI:	<he hecho la comunión> [/] he hecho la &comu comunión (..) Jua:n [?]

(.) mi pima: [*] e:h mi mano [*] (.) el Jaime (.) el Iván (.) el

Rafa y: quién más [=! deleted /s/, dialectal]

[=! looks for the help of MOT, looking in her direction although

she is not in the room]?

%err:	pima=prima $PHO mano=hermano $PHO

%xepr:	$i5:MMN:REP; $et3:DRA:DIS; $i5:MMN:ORD:OMI; $et3:DRA:FIL;

*CHI:	eh@i (.) quién má [=! deleted /s/, dialectal] [=! shakes head no] ?

%xepr:	$et3:DRA:ERQ; $et3:GSA

*MOT:	venga@i Ana cuéntaselo todo .

*CHI:	el Javiér [=! deleted /r/, dialectal] (.) e:h Mario (...) y: (..) <me

regalaron> [=! very loose articulation] <muchas cosas>

[=! deleted /s/ in final position of both words, dialectal] (...).

%xepr:	$et3:DRA:FIL

*CHI:	y: [/] (0.6) y: (...) ya'ta@d .

*MOT:	ya'ta:@d ?

*CHI:	sí .

*MOT:	y: dónde [/] dónde estuvisteis

[=! aspirated /s/ in postnuclear position of the syllable,

deleted /s/ in final position of the word, both dialectal] ?

*CHI:	a: [*] Carcola [*].

%err:	a=en la $MOR $PHO; Carcola=Caracola $PHO

%xepr:	$i5:MMN:ORD:OMI

*MOT:	a:h@i y eso dónde está [=! aspirated /s/, dialectal] ?

*CHI:	está [=! aspirated /s/, dialectal] a:[*] Virgen .

%err:	a=en la $MOR $PHO

*MOT:	en la Virgen ?

*CHI:	al la:o [=! deleted /d/ between vocals, dialectal].

*MOT:	al lao no .

*CHI:	sí: .

*MOT:	na:h (...) y qué <más cositas>

[=! deleted /s/ in final position of both words, dialectal] a+ver@i

dile algo más [=! deleted /s/, dialectal] .

*CHI:	(0.3) [=! sighs] hum@i (0.5) que e [*] digo ?

%err:	e=le $PHO

%xepr:	$i5:MQT:RVP:INT

*MOT:	xxx xxx xxx xxx xxx ?

*CHI:	ya l'he dicho .

*MOT:	ah@i ya se l'has [=! deleted /s/, dialectal] dicho [>] ?

*BOY:	no: [<].

*CHI:	sí [=! looking at BOY] !

*BOY:	has [=! deleted /s/, dialectal] dicho +"/.

*BOY:	+" <muchos regalos> [?] [=! deleted /s/, dialectal] .

*BOY:	pero no l'has [=! deleted /s/, dialectal] dicho <lo que xxx xxx> [>].

*MOT:	no l'has [=! deleted /s/, dialectal] dicho [<] lo [/-] qué regalos.

*CHI:	e:h u:n jama [*]

tablet [=! deleted /t/ at the end of the word, dialectal] xxx xxx

chanclas [=! deleted /s/, dialectal] artera [*] estuche [*] el boli

xxx xxx [=! counting with fingers] +/.

%err:	jama=pijama $PHO; tablet=una tablet $MOR; chanclas=unas chanclas

$MOR; artera=una cartera $PHO $MOR; estuche=un estuche $MOR

%xepr:	$et3:DRA:FIL; $i5:MMN:ORD:OMI; $i5:MMN:ORD:OMI;

$i5:MMN:ORD:OMI; $i5:MMN:ORD:OMI

*MOT:	y cuál es [=! deleted /s/, dialectal] el que más

[=! deleted /s/, dialectal] t'ha gustao

[=! deleted /d/ between vocals, dialectal] ?

*CHI:	hum@i [=! laughs] .

%xepr:	$i5:MQT:RVP:INT

*MOT:	eh@i ?

*CHI:	0.

*MOT:	cuál es [=! deleted /s/, dialectal] el que te [/-] más

[=! deleted /s/, dialectal] t'ha gustao

[=! deleted /d/ between vocals, dialectal] ?

*CHI:	a [*] tablet [=! deleted /t/ at the end of the word, dialectal] .

%err:	a=la $PHO

*MOT:	la table: ?

*CHI:	sí [=! smiles ].

@End
