## Supplemental file 1 for "Language impairment with a microduplication in 1q42.3q43"

**Inventory for Client and Agency Planning (ICAP)**

| **Escalas de conducta adaptativa (a)** | Punt.  Escala (b) | Age Equivalent  © | EE-EC | Rango Instructivo  (d) | Punt.  Difer  Escala |
| --- | --- | --- | --- | --- | --- |
| Motor Skills | 477 | 4 -9 | -4 - 11 | 3-5 a 6-0 | PER(e):<1  PT (f): 52  IRR (g): 45/90 |
| Social and Communication Skills | 487 | 5-1 | -2 - 9 | 3-0 a 9-0 | PER:9  PT: 80  IRR: 73/90 |
| Personal Living Skills | 500 | 9-11 | +3 meses | 7-7 a 13-1 | PER:53  PT: 101  IRR: 91/90 |
| Community Living Skills | 483 | 7-3 | -2 - 5 | 5-9 a 8-11 | PER:10  PT: 81  IRR: 71/90 |
| Broad Independence (Total) | 487 | 6-11 | -2 - 9 | 4-11 a 9-7 | PER:9  PT: 80  IRR: 75/90 |
| Scoring: SS (Standard Score); PR (Percentile Rank); AE (Age Equivalent) | | | | | |

**Illinois Test of Psycholinguistic Abilities (ITPA)**

|  | NIVEL REPRESENTATIVO | | | | | | NIVEL AUTOMÁTICO | | | | |  |
| --- | --- | --- | --- | --- | --- | --- | --- | --- | --- | --- | --- | --- |
|  | Comprensión | | Asociación | | Expresión | | Integración | | Mem secuencial | | T.complementario |  |
|  | Audit | Visual | Audit | Visual | verbal | Moto | Gram | Visual | Audit | Vmot | Int. audit |  |
| 52 | **0(24)** | **0(28)** | **0(24)** | **0(28)** | **0(24)** | **0(26)** | **0(25)** | **0(34)** | **0(24)** | **0(34)** | **0(26)** | 52 |
| 50 |  |  |  |  |  |  |  |  |  |  |  | 50 |
| 48 |  |  |  |  |  |  |  |  |  |  |  | 48 |
| 46 |  |  |  |  |  |  |  |  |  |  |  | 46 |
| 44 |  |  |  |  |  |  |  |  |  |  |  | 44 |
| 42 |  |  |  |  |  |  |  |  |  |  |  | 42 |
| 40 |  |  |  |  |  |  |  |  |  |  |  | 40 |
| 38 |  |  |  |  |  |  |  |  |  |  |  | 38 |
| 36 |  |  |  |  |  |  |  |  |  |  |  | 36 |
| 34 |  |  |  |  |  |  |  |  |  |  |  | 34 |
| 32 |  |  |  |  |  |  |  |  |  |  |  | 32 |
| 30 |  |  |  |  |  |  |  |  |  |  |  | 30 |
| 28 |  |  |  |  |  |  |  |  |  |  |  | 28 |
| 26 |  |  |  |  |  |  |  |  |  |  |  | 26 |
| 24 |  |  |  |  |  |  |  |  |  |  |  | 24 |
| 22 |  |  |  |  |  |  |  |  |  |  |  | 22 |
| 20 |  |  |  |  |  |  |  |  |  |  |  | 20 |

**Registro Fonológico Inducido [Induced Phonological Register] (RFI)**

| ITEMS | Spontaneous expression | **Listen and repeat** |
| --- | --- | --- |
| 1. moto | + |  |
| 2. boca | + |  |
| 3. piña | + |  |
| 4. piano |  | + |
| 5. pala | + |  |
| 6. pie | + |  |
| 7. niño | + |  |
| 8. pan | + |  |
| 9. ojo | + |  |
| 10.llave | + |  |
| 11.luna | + |  |
| 12.campana | capama |  |
| 13.indio | idio |  |
| 14.toalla |  | + |
| 15.fuma | + |  |
| 16.dedo | deo |  |
| 17.peine | peme |  |
| 18.ducha |  | + |
| 19.gafas | + |  |
| 20.toro | + |  |
| 21.silla | + |  |
| 22.taza | zaza |  |
| 23.cuchara |  | Chuchara |
| 24.teléfono |  | Leno |
| 25.sol | + |  |
| 26.casa | + |  |
| 27.pez | + |  |
| 28.jaula |  | + |
| 29.zapato |  | + |
| 30.flan | fran |  |
| 31.lápiz | + |  |
| 32.pistola | pitola |  |
| 33.mar |  | + |
| 34.caramelo | camelo |  |
| 35.plátano | plano |  |
| 36.globo | golo |  |
| 37.palmera | paera |  |
| 38.clavo | cavo |  |
| 39.tortuga | tuga |  |
| 40.pueblo | porvo |  |
| 41.tambor | pambor |  |
| 42.escoba | coba |  |
| 43.mariposa | posa |  |
| 44.puerta | + |  |
| 45.bruja | + |  |
| 46.grifo | + |  |
| 47.jarra | jaja |  |
| 48.tren | + |  |
| 49.gorro | goro |  |
| 50.rata | rato |  |
| 51.cabra | paca |  |
| 52.lavadora | vadora |  |
| 53.preso | peso |  |
| 54.semáforo | famo |  |
| 55.fresa | fesa |  |
| 56.árbol | abo |  |
| 57.periódico | pediodo |  |

| observation |
| --- |
| + Not failures in output the word  The girl shows shame because she is fully aware of her problem and knows that she will not pronounce it correctly, so she shows behaviors such as hiding her face, nervous laughter and takes a long time to utter the word. When she dominates the word he emits it quickly. |

**Batería para la Evaluación de los Procesos Lectores en Secundaria y Bachillerato [Battery for the Evaluation of Reading Abilities in High School Students] (PROLEC-SE)**

| **PROCESOS LEXICOS** | **PUNTUACIÓN DIRECTA** | | **PUNTUACIÓN CENTIL** | |
| --- | --- | --- | --- | --- |
|  | madre | padre | madre | padre |
| Palabras | 40 | 40 | 95 | 95 |
| Pseudopalabras | 40 | 40 | 95 | 95 |
| **PROCESOS SINTÁCTICOS** | **PD** | | **PC** | |
| Emparejamiento  dibujo-oración | 24  Sin dificultad NIVEL ALTO | 24  Sin dificultad  NIVEL ALTO |  |  |
| Signos de puntuación | 24  Sin dificultad  NIVEL ALTO | 24  Sin dificultad  NIVEL ALTO | 90 | 90 |
| **PROCESOS SEMÁNTICOS** | **PD** | | **PC** | |
| Comprensión de textos | 14  Sin dificultad  NIVEL MEDIO | 13  Sin dificultad  NIVEL BAJO | 45 | 25 |
| Estructura de texto | 17  Sin dificultad  NIVEL BAJO | 13  Sin dificultad  NIVEL BAJO | 25 | <5 |
| **TOTAL DE LA BATERÍA** | **PD** | | **PC** | |
|  | **158**  **Sin dificultad**  **NIVEL MEDIO** | **138**  **Existe dificultad** | **75** | **<5** |

**Test de Aptitud Verbal "Buenos Aires" [Buenos Aires Verbal Aptitude Test] (BAIRES)**

|  | Puntuación Directa | | PT (*puntuación típica*) | |
| --- | --- | --- | --- | --- |
|  | madre | padre | madre | padre |
| Definiciones | 14 | 13 | 47 | 43 |
| Sinónimos | 8 | 6 | 43 | 39 |
| Total (Def. + Sin.) | 22 | 19 | 90 | 82 |

**WAIS-III**

PADRE: pruebas verbales

|  | PRUEBAS VERBALES | | | | | |  | Coeficiente Intelectual verbal | | |
| --- | --- | --- | --- | --- | --- | --- | --- | --- | --- | --- |
|  | Información | Comprensión | Aritmética | Semejanzas | Dígitos | Vocabulario |  |  |  |  |
| 20 | 0 | 0 | 0 | 0 | 0 | 0 | 20 | --- | -- | -- |
| 19 | 0 | 0 | 0 | 0 | 0 | 0 | 19 | 145 | SUPERIOR | |
| 18 | 0 | 0 | 0 | 0 | 0 | 0 | 18 | 140 |  |  |
| 17 | 0 | 0 | 0 | 0 | 0 | 0 | 17 | 135 |  |  |
| 16 | 0 | 0 | 0 | 0 | 0 | 0 | 16 | 130 |  |  |
| 15 | 0 | 0 | 0 | 0 | 0 | 0 | 15 | 125 |  |  |
| 14 | 0 | 0 | 0 | 0 | 0 | 0 | 14 | 120 |  |  |
| 13 | 0 | 0 | 0 | 0 | 0 | 0 | 13 | 115 | MEDIO | |
| 12 | 0 | 0 | 0 | 0 | 0 | 0 | 12 | 110 |  |  |
| 11 | **x** | 0 | 0 | **x** | 0 | 0 | 11 | 105 |  |  |
| 10 | 0 | **x** | **x** | 0 | **x** | **x** | 10 | **100** |  |  |
| 9 | 0 | 0 | 0 | 0 | 0 | 0 | 9 | 95 |  |  |
| 8 | 0 | 0 | 0 | 0 | 0 | 0 | 8 | 90 |  |  |
| 7 | 0 | 0 | 0 | 0 | 0 | 0 | 7 | 85 |  |  |
| 6 | 0 | 0 | 0 | 0 | 0 | 0 | 6 | 80 | INFERIOR | |
| 5 | 0 | 0 | 0 | 0 | 0 | 0 | 5 | 75 |  |  |
| 4 | 0 | 0 | 0 | 0 | 0 | 0 | 4 | 70 |  |  |
| 3 | 0 | 0 | 0 | 0 | 0 | 0 | 3 | 65 |  |  |
| 2 | 0 | 0 | 0 | 0 | 0 | 0 | 2 | 60 |  |  |
| 1 | 0 | 0 | 0 | 0 | 0 | 0 | 1 | 55 |  |  |
| 0 | 0 | 0 | 0 | 0 | 0 | 0 | 0 | --- |  |  |

MADRE: pruebas verbales

|  | PRUEBAS VERBALES | | | | | |  | Cociente verbal | |
| --- | --- | --- | --- | --- | --- | --- | --- | --- | --- |
|  | Información | Comprensión | Aritmética | Semejanzas | Dígitos | Vocabulario |  |  |  |
| 19 | 0 | 0 | 0 | 0 | 0 | 0 | 19 | 145 | SUPERIOR |
| 18 | 0 | 0 | 0 | 0 | 0 | 0 | 18 | 140 |  |
| 17 | 0 | 0 | 0 | 0 | **x** | 0 | 17 | 135 |  |
| 16 | 0 | 0 | 0 | 0 | 0 | 0 | 16 | 130 |  |
| 15 | 0 | 0 | 0 | 0 | 0 | 0 | 15 | 125 |  |
| 14 | 0 | 0 | 0 | 0 | 0 | 0 | 14 | 120 |  |
| 13 | 0 | 0 | 0 | 0 | 0 | 0 | 13 | 115 | MEDIO |
| 12 | 0 | 0 | 0 | **x** | 0 | 0 | 12 | 110 |  |
| 11 | 0 | 0 | 0 | 0 | 0 | **x** | 11 | **105** |  |
| 10 | **x** | 0 | 0 | 0 | 0 | 0 | 10 | 100 |  |
| 9 | 0 | **x** | 0 | 0 | 0 | 0 | 9 | 95 |  |
| 8 | 0 | 0 | x | 0 | 0 | 0 | 8 | 90 |  |
| 7 | 0 | 0 | 0 | 0 | 0 | 0 | 7 | 85 |  |
| 6 | 0 | 0 | 0 | 0 | 0 | 0 | 6 | 80 | INFERIOR |
| 5 | 0 | 0 | 0 | 0 | 0 | 0 | 5 | 75 |  |
| 4 | 0 | 0 | 0 | 0 | 0 | 0 | 4 | 70 |  |
| 3 | 0 | 0 | 0 | 0 | 0 | 0 | 3 | 65 |  |
| 2 | 0 | 0 | 0 | 0 | 0 | 0 | 2 | 60 |  |
| 1 | 0 | 0 | 0 | 0 | 0 | 0 | 1 | 55 |  |
| 0 | 0 | 0 | 0 | 0 | 0 | 0 | 0 | ---- |  |
